# Mechanism of hyperproteinemia-induced blood cell homeostasis imbalance in an animal model

**DOI:** 10.1101/2021.08.09.455622

**Authors:** Guang Wang, Yongfeng Wang, Jianglan Li, Ruji Peng, Xinyin Liang, Xuedong Chen, Guihua Jiang, Jinfang Shi, Yanghu Si-ma, Shiqing Xu

## Abstract

Hyperproteinemia is a metabolic disorder associated with increased plasma protein concentration (PPC). It is often clinically complicated by malignant diseases or severe infections. Research on the molecular mechanism of High PPC (HPPC) is scant. Here, an animal model of primary hyperproteinemia was constructed in an invertebrate, *Bombyx mori*, to investigate the effect of HPPC on circulating blood cells. We showed that HPPC affected blood cell homeostasis and enhanced blood cell phagocytosis, leading to increased reactive oxygen species levels, and induced programmed cell death that depended on the endoplasmic reticulum-calcium ion signaling pathway. HPPC induced the proliferation of blood cells, mainly granulocytes, by activating the JAK/STAT signaling pathway. Supplementation with endocrine hormone active substance 20E significantly reduced the impact of HPPC on blood cell homeostasis. Herein, we reported a novel signaling pathway by which HPPC affected blood cell homeostasis, which was different from hyperglycemia, hyperlipidemia, and hypercholesterolemia. In addition, we showed that down-regulation of gene expression of the hematopoietic factor *Gcm* could be used as a potential early monitoring index for hyperproteinemia.

**Highlights:**

• HPPC induced proliferation of circulating blood cells through the JAK/STAT pathway, leading to a significant increase in the proportion of granulocytes.
• Supplementing endocrine hormone 20E improved hematopoietic function and restored the homeostasis of circulating blood cells through modulation of the JAK/STAT signaling pathway.
• HPPC induced PCD in blood cells through the endoplasmic reticulum-calcium ion release signaling pathway.
• Down-regulation of the gene expression of the hematopoietic factor *Gcm* could be used as a potential early monitoring index for hyperproteinemia.

## Introduction

Hyperproteinemia is a metabolic disorder with persistently abnormally elevated plasma protein concentration (PPC), which often clinically complicates multiple myeloma (***Hussain et al., 2019; Chang et al., 2019***), nephropathy (***Oliveira et al., 2019; Kitazawa et al., 2018***), liver cirrhosis (***Fujii et al., 2014***), infection (***Kluck et al., 2019***), metabolic alkalosis (***Matoušek et al., 2018***), pneumonia (***Gerin et al., 2014***), and other serious diseases, adversely affecting the prognosis of patients (***Abuzaid et al., 2020; Hussain et al., 2019***). It also appears after plasma therapy or treatment with intravenous immunoglobulin (***Boyle et al., 2011; Steinberger et al., 2003***). Because it is clinically difficult to distinguish between hyperproteinemia and the primary disease or the impact of infection on the body, there is no reliable primary disease model of hyperproteinemia (***Chen et al., 2018; Riemer et al., 2016***). Therefore, there is a paucity of research on the pathological mechanism of hyperproteinemia, and research on the effect of hyperproteinemia on the hematological system is scant.

A large number of studies have shown that blood system metabolic disorders, such as hyperglycemia, hyperlipidemia, and hypercholesterolemia are associated with chronic inflammation (***Wagner et al., 2021; Zhao et al., 2020***), which destroys the homeostasis of blood cells leading to severe disproportion of circulating blood cells; monocytes and neutrophils increase significantly (***Aguilar-Ballester et al., 2020; Nagareddy et al., 2013***); it also affects the proliferation and differentiation of blood cells (***Gu et al., 2019; Barrett et al., 2019***). It is worth noting that patients with hyperproteinemia also commonly experience abnormal metabolic changes, such as deranged blood glucose and blood lipid levels (***Kluck et al., 2019; Hussain et al., 2019***), suggesting, thereby that hyperproteinemia may have a complex effect on blood homeostasis. It has also been clinically found that many diseases with associated hyperproteinemia show a disproportion in the types of circulating blood cells (***Sant et al., 2020; Kitazawa et al., 2018***), such as increased neutrophils in patients with multiple myeloma (***Hussain et al., 2019***); patients with metabolic syndrome have an increase in the number of non-classical monocytes (***Khan et al., 2016***). Thus, it is necessary to investigate the effect of primary high PPC (HPPC) on blood cell homeostasis.

Although hyperproteinemia is commonly observed in the clinic (***Kitazawa et al., 2018; Bergstedt and Lingen, 1957***), it has not been possible to use animal models, such as mammals and fruit flies to establish primary hyperproteinemia disease models (***Chen et al., 2018; Riemer et al., 2016***). We selected an invertebrate, *Bombyx mori* (***He et al., 2019; Tabunoki et al., 2016***), to construct an animal model of HPPC (AM) with no primary disease impact and controllable PPC levels; we have previously found that HPPC had a complex effect on the development of gonads and remodeling of the fat composition of the metabolic tissue (***Wang et al., 2019; Chen et al., 2018***). We further observed previously that HPPC affected the expression of antimicrobial peptides by activating the Toll pathway and Imd pathway of NF-κB signaling, showing inconsistent changes in antimicrobial activity against Gram-negative and -positive bacteria. Furthermore, HPPC inhibited phenoloxidase synthesis and weakened the immune effect of melanin in hemolymph (***Wang et al., 2021***). Silkworm has a completely open circulating hemolymph system, and hematopoiesis depends on the proliferation and differentiation of blood cells in hematopoietic organs and circulating hemolymph (***Nakahara et al., 2010; Ling et al., 2005***). In the silkworm model of hyperproteinemia, there are significant changes in number of the blood cells (***Wang et al., 2021***), but the relevant mechanism is still unclear. To this end, this study aimed to elucidate the mechanism of HPPC’s influence on the proliferation and differentiation (hematopoiesis) of circulating blood cells.

## Results

### HPPC induced changes in blood cell homeostasis

AO-PI staining can accurately identify the five types of hemocytes (blood cells) of the silkworm: Plasmatocyte (Pla), Prohemocyte (Pro), Granulocyte (Gra), Oenocytoid (Oen), and Spherulocyte (Sph) (***Figure 1A***). Compared with the CK group, HPPC caused an increase in blood cell density in the early and mid-term period (48 h to 96 h) after modeling, and a significant decrease in the later period (144 h to 192 h) in the AM (***Figure 1B and Figure 1—figure supplement 1B***). With a significant increase in PPC starting 48 h after modeling, the density of the model group (AM) Pla continued to decrease until 144 h when there was only a trace amount; the density of Pro, Gra, Oen, and Sph all showed a similar trend of first increasing and then decreasing. Among them, the density of Sph dropped rapidly from 96 h to trace amounts, while the densities of Pro, Gra, and Oen were significantly higher than the control group before 96 h, and thereafter gradually decreased. Both Pro and Oen decreased to a low density, significantly lower than the control group at 144 h, while Gra maintained a similar density to CK after 144 h (***Figure 1—figure supplement 1C-G***). These results showed that HPPC had the greatest adverse effect on the density of Pla.

**Figure 1.**
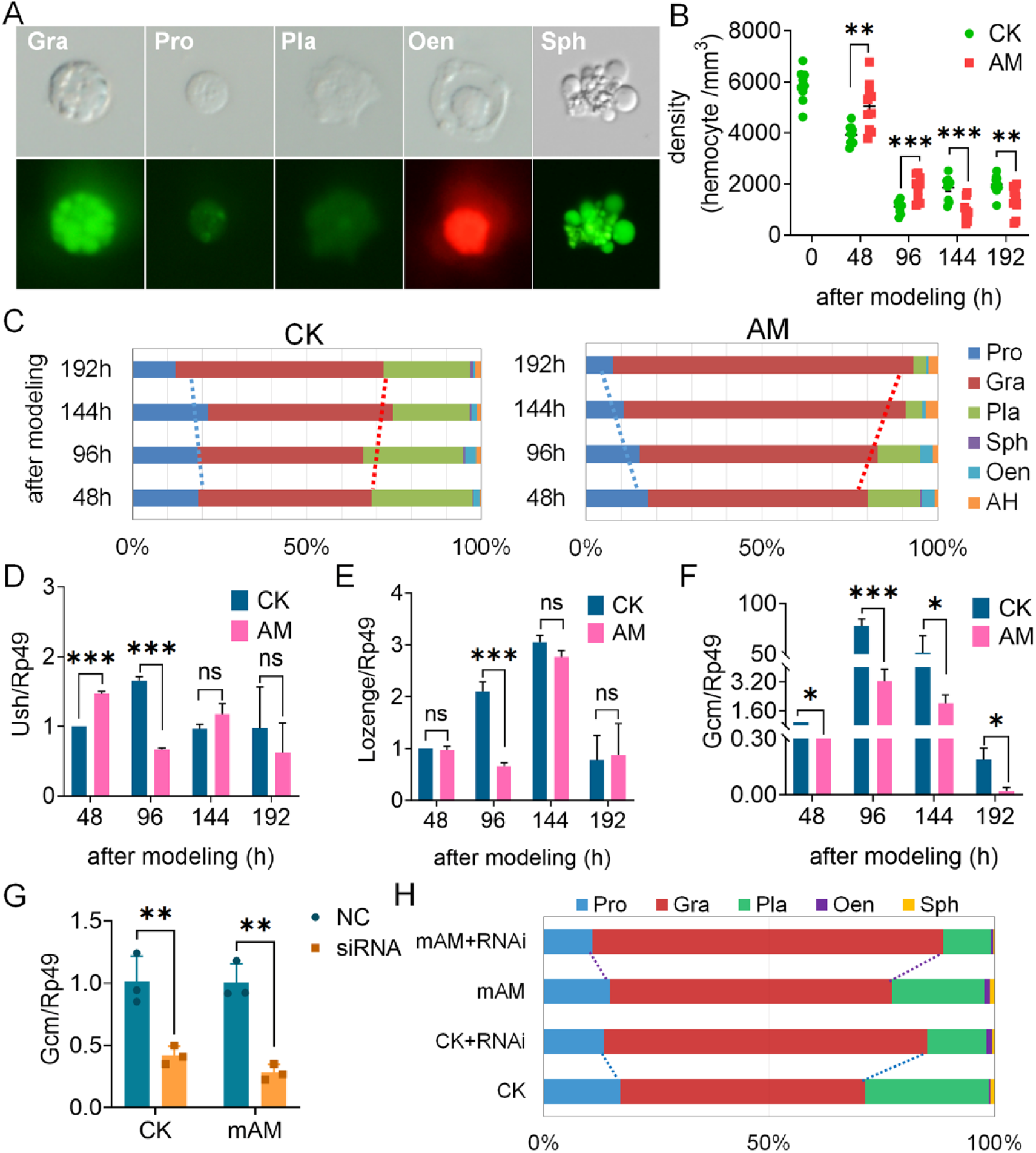
HPPC affected the composition of circulating hemocytes (blood cells). (**A**) Results of blood cell classification after AO-PI staining. Pro, Prohemocyte. Pla, Plasmatocyte. Gra, Granulocyte. Sph, Spherulocyte. These four types of hemocytes were all stained with green fluorescence by AO, which could be distinguished from the intensity of fluorescence and the morphology of hemocytes. Oen, Oenocytoid, which could not be stained by AO, but was stained with red fluorescence by PI. The blood cell nucleus with necrosis or membrane rupture could also be stained red by PI, which could be distinguished from the other four types of hemocytes in morphology. (**B**) Blood cell density (n = 10). CK, control group; AM, model group. (**C**) Percentage composition of circulating hemocytes (n = 3). AH, could not distinguish the types, often deformities. (**D-F**) Gene transcription levels of hematopoietic-related factors (n = 3). D, *Ush*; E, *Lozenge*; F, *Gcm*. The internal reference gene was *Bombyx mori* Rp49. (**G-H**) Interference with *Gcm* gene at the individual level affected blood cell composition. The *Gcm* gene transcription level of hemocytes (**G**) and the percentage composition of blood cell classification (**H**) were investigated 48 h after the intervention treatment (n = 3). CK, control group. CK+RNAi, individual intervention in CK group. mAM, mild model group. mAM+RNAi, individual intervention in the mild model group. Each silkworm in the siRNA group was injected with 10 μg Gcm-siRNA, the control (NC in Figure 1G, and CK or mAM in Figure 1H) was injected with the same amount of negative control siRNA (si-NC), and each group was effectively injected with 18 individuals (6 individualsÍ3 repetitions). Data are shown as mean ± SEM. ns p>0.05,*p<0.05, **p<0.01, and ***p<0.001 by Student’s T-test. The online version of this article includes the following source data and figure supplement(s) for figure 1: **Source data 1.** Source data for ***Figure 1B, Figure 1C, Figure 1H, Figure 1—figure supplement 1.*** **Figure supplement 1.** Changes of plasma protein concentration (PPC) after modeling. **Figure supplement 2.** Impact of HPPC on development and surviving rate of Bombyx mori **Figure supplement 3.** The effect of hyperproteinemia on the density of different types of blood cells.

Percentage composition (PC) of the five types of hemocytes showed that the homeostasis of blood cells in the hemolymph of the AM group was severely deranged (***Figure 1C***). From 0 h to 192 h after modeling, compared with the CK group, the PC value of Gra in the AM group continued to increase with time after modeling, reaching up to 62.38% and 85.50% at 48 h and 192 h, respectively, 12.63% and 25.95% higher than the control, respectively. The high PC value of Pro and Pla at the beginning of modeling decreased rapidly with the continuous increase of Gra, from 18.87% and 28.92% at 48 h to 7.63% and 3.63% at 192 h, respectively. Sph were almost invisible 96 h after modeling. Thus, it can be seen that Gra was the most abundant blood cell type in the hemolymph of the hyperproteinemia silkworm model, and it was also an indicator blood cell that changed significantly under the influence of HPPC.

The transcriptional level of genes of potential hematopoietic-related factors in blood cells has been shown to be significantly affected by HPPC. Compared with the CK group, the *Ush* gene and the *Lozenge* gene that are related to Oen production in the AM group were up-regulated at 48 h after modeling, and both genes were significantly down-regulated at 96 h; thereafter, these gene levels returned to the level of the CK group at 144 h (***Figure 1D-E***). It is worth noting that the transcriptional level of homologous gene *Gcm* in the silkworm, which is the *Drosophila* Pla-promoting factor, showed a cliff-like down-regulation after modeling compared with the control group (***Figure 1F***). This suggested that HPPC may change the ability of blood cell formation by modulating the expression of hematopoietic-related factors, such as *Gcm*.

The pathological effect of the AM group was serious, and all the individuals died. In order to explore the role of hematopoietic factor expression in the development of hyperproteinemia, a silkworm model of hyperproteinemia was introduced, which can complete the individual development and has less pathological effect. Moreover, in order to explore the relationship between *Gcm* gene expression level and the changes of hemocyte composition in hyperalbuminemia silkworm, a mild disease model was introduced (***Figure 1—figure supplement 1A***).

After mAM modeling, *Gcm* gene siRNA was injected immediately; 48 h later, it was found that the *Gcm* gene mRNA levels of blood cells in the CK and mAM groups were down-regulated by 59.37% and 72.51%, respectively (***Figure 1G and Figure 1— figure supplement 1A***), and the PC value of Pla decreased by 14.33% (CK) and 9.89% (mAM); the Pro PC value decreased by 3.57% (CK) and 3.96% (mAM); while Gra PC value increased by 17.58% (CK) and 15.21% (mAM) (***Figure 1H***). This showed that by silencing the *Gcm* gene, not only did the CK group show a similar blood cell homeostasis trend as the AM group, but the blood cell homeostasis changed in the mAM group also showed the superimposed effect of the increase of PPC and the interference of *Gcm* gene expression. It further showed that the influence of HPPC on the regulation of the production of different types of blood cells was mutually constrained.

### HPPC affects blood cell proliferation and differentiation through the JAK/STAT pathway

The foregoing results showed that HPPC led to a decrease in the number of hemocytes and changes in the homeostasis of circulating blood cells. In order to explain this phenomenon, the changes in the proliferation and differentiation of blood cells were first investigated

The proliferation level of circulating hemocytes investigated by Edu staining, showed that the AM group had more active cell proliferation in the later stage. At 48 h after modeling, there was no significant difference in the Edu-positivity rates of hemocytes in the CK group and the AM group, which were 2.32% and 2.11%, respectively, of the base physiological level. The Edu-positivity rate of the CK group increased to 7.84% at 96 h, was 6.62% at 144 h, and was restored to the base level of 2.12% at 192 h. Nevertheless, the Edu-positivity rate in AM group increased significantly from 2.49% at 96 h to 8.88% at 144 h, and remained at a high level of 8.27% at 192 h (***Figure 2A-B***). In the time period, 96 h to 144 h after modeling in the CK group, an active proliferation period, the Edu-positive hemocytes were mainly granulocytes with a small number of prohemocytes; while in the AM group, only granulocytes were observed in the Edu-positive cells from 144 h to 192 h (***Figure 2C***). The results of repeated Edu staining investigations were consistent with these results (***Figure 2—figure supplement 2E-F***). This implied that the proliferation of circulating hemocytes in the AM group was delayed compared to that in the CK group, but in the later stages after modeling, active hemocyte proliferation continued to occur, with the type of blood cell being mainly granulocytes.

**Figure 2.**
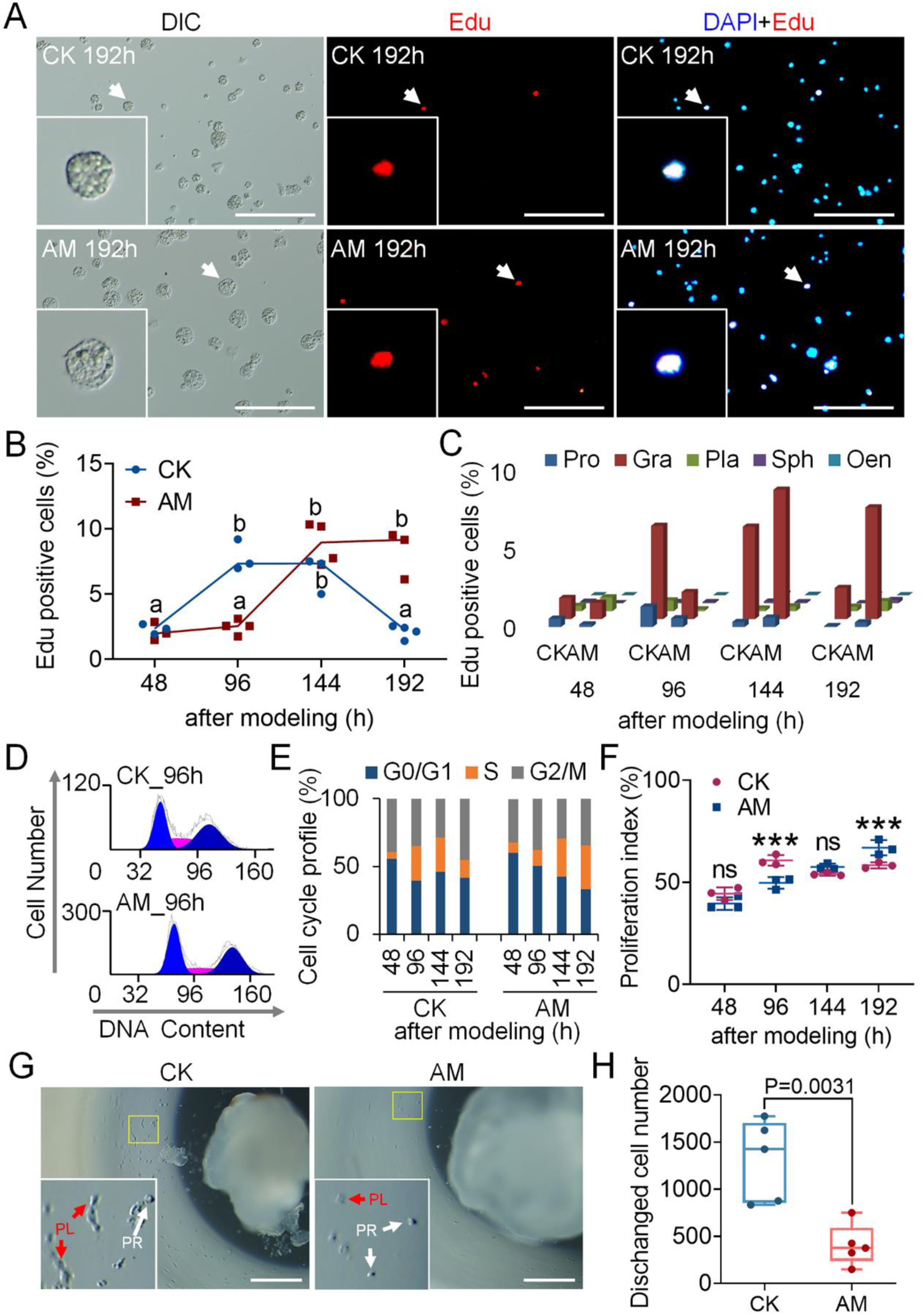
Hyperproteinemia led to delay proliferation of hemocytes in the later stages of modeling. (**A-C**) DNA staining of proliferating hemocytes. Edu staining and DAPI staining respectively marked the proliferating cell nuclei and all cell nuclei in the hemolymph at 48 h, 96 h, 144 h, and 192 h after modeling. Positive rate (%) = (Edu-positive cell number / DAPI-positive cell number) × 100. Scale bars 100 μm. (**A**) Fluorescence of Edu staining at 192 h after modeling. (**B**) Edu-positive rate of hemocytes (CK 48 h – 192 h, n = 3, 3, 3, 4; AM 48 h – 192 h, n = 3, 4, 4, 3. P≤0.05). (**C**) Edu-positive rate of different types of hemocytes. There were significant differences between groups with different letters. (**D**) PI-FCM was detected the cell cycle phase of circulating hemocytes (n = 3). The cell cycle was divided into G0/G1, S and G2/M phases. (**E**) Quantitative diagram of the phase of cell cycle. The ordinate indicated the proportion of cells in the G0/G1 phase, S phase and G2/M phase to the total number of cells. (**F**) Proliferation index (PI) (n = 3). It indicated the proportion of proliferating hemocytes to the total number of circulating hemocytes. PI (%) = (S+G2/M) / (G0/G1+S+G2/M) × 100. Data are shown as mean ± SEM. ns p>0.05,*p<0.05, **p<0.01, and ***p<0.001 by Student’s T-test. (**G-H**) Culture HPO-Wing organize complex *in vitro*. After 24 hours of modeling, the HPO-Wing complex was taken out, and the level of hematopoiesis was investigated after 48 hours of culture *in vitro*. PR, Prohemocyte. PL, Plasmatocyte. (**G**) HPO-Wing morphology. Scale bars 0.5 mm; (**H**) The number of hemocytes released by a single culture of HPO-Wing (n = 5). Data are shown as mean ± SEM. P-values of significance are calculated by Student’s T-test. The online version of this article includes the following source data and figure supplement(s) for figure 2: **Source data 1.** Source data for ***Figure 2B, Figure 2C, Figure 2E, Figure 2F, Figure 2H, Figure 2—figure supplement 2B, Figure 2—figure supplement 2C and Figure 2—figure supplement 2D.*** **Figure supplement 1.** HPPC affected the cycle phase distribution of circulating hemocytes. **Figure supplement 2.** Hyperproteinemia led to arrest proliferation of hemocytes at 96h after modeling.

In order to explore the cause of delayed proliferation of circulating hemocytes observed in HPPC, the cell cycle phase of hemocytes was investigated (***Figure 2D and Figure 2—figure supplement 2A***). At 48 h to192 h after modeling, the proportion of hemocytes in the G0/G1 phase and S phase in the AM group showed a trend opposite to that of the CK group (***Figure 2—figure supplement 2B and D***). At 96 h after modeling, the cells were arrested in the G0/G1 phase, indicating that HPPC inhibited DNA synthesis of hemocytes (***Figure 2E***). The cell proliferation index also showed a significant difference at 96 h and 192 h after modeling, and converse changes between the CK group and the AM group were observed (***Figure 2E-F***). This showed that the distribution of blood cells in the cell cycle had changed during the course of hyperproteinemia. In the early stages after modeling, hemocytes were arrested in the G1 stage, but vigorous cell division in the later stages continued, thereby displaying the phenomenon of delayed cell proliferation.

To assess the effect of HPPC on hematopoietic capacity, a previously established method of *in vitro* culture of hematopoietic organs was used. The HPO-Wing organize complex was taken out 24 h after modeling, and the level of hematopoiesis was observed after 48 h of *in vitro* culture. In the CK group, HPO-Wing was at the initial stage of normal dissociation of development by metamorphosis. Although the tissues were shrinking, they could still proliferate and release a large number of mature hemocytes, including prohemocytes and plasmatocytes (***Figure 2G***). In the HPO-Wing of the AM group, however, although the morphology was intact and the tissues had not begun to dissociate, the number of released hemocytes was very small (***Figure 2G-H***). This implied that even if the HPO left the HPPC environment *in vivo*, its hematopoietic ability was still severely adversely affected by HPPC in the body during the initial 24 h of modeling.

To study the molecular signaling pathways by which HPPC affected the proliferation of hemocytes, we focused on the JAK/STAT pathway. Immunofluorescence was used to detect the STAT protein in hemocytes. In both the CK group and the AM group, there were STAT-positive hemocytes (***Figure 3A***). At 48 h to 96 h after modeling, there was no significant difference in the STAT-positivity rate of CK and AM hemocytes, which increased from 1.40% and 1.28% to 3.72% and 3.18%, respectively. The STAT-positivity rate of the CK group fell back to 1.43% at 192 h; meanwhile the rate of AM group significantly increased to 8.11% (***Figure 3B***). Classification and statistical analysis of STAT-positive hemocytes showed that the main cell type was the granulocyte (***Figure 3—figure supplement 3C***). The changes in protein content by western blotting also showed that STAT in the AM group was no significant difference in the CK group at the early and middle stages (48 h and 96 h) after modeling, but in the late stage (192 h) was significantly up-regulated (***Figure 3C and Figure 3—figure supplement 3D***).

**Figure 3.**
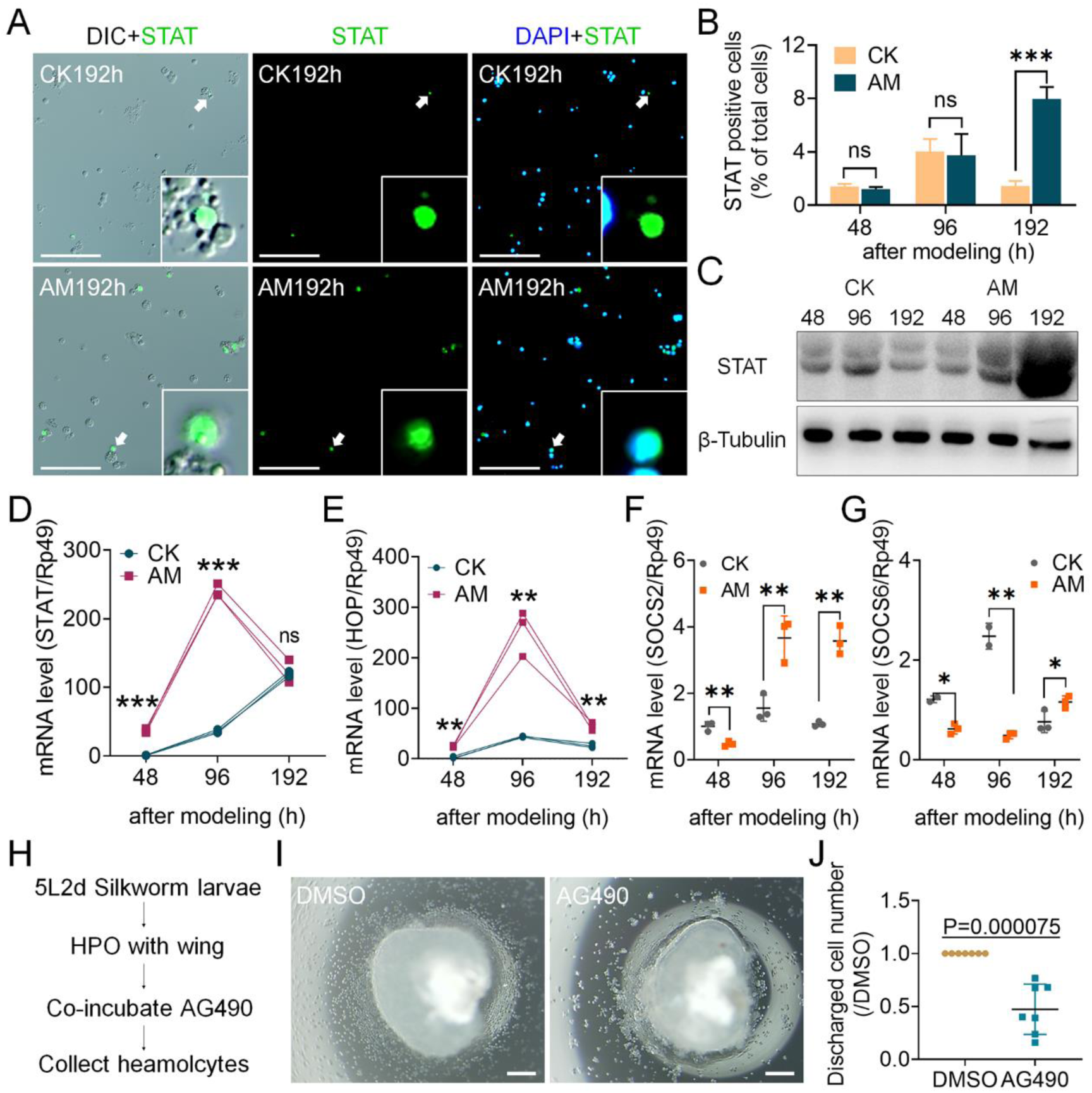
HPPC activated the JAK/STAT signal pathway. STAT immunofluorescence and DAPI staining marked the cells expressing STAT protein and all the cell nuclei in the hemolymph at 48 h, 96 h and 192 h after modeling, respectively. (**A**) STAT immunofluorescence image of hemocytes at 192 h after modeling. Scale bars 100 μm. (**B**) STAT-positive rate of hemocytes (n = 3). STAT positive rate (%) = (STAT positive cell number / total cell number) × 100. (**C**) The STAT protein in blood cell was analyzed by western blotting and referenced by *β-Tubulin* (n = 3). (**D-G**) qPCR analyzed the transcription level of JAK/STAT pathway genes. The reference gene was *Rp49* (n = 3). Data are shown as mean ± SEM. ns p>0.05,*p<0.05, **p<0.01, and ***p<0.001 by Student’s T-test. (**H-J**) Culture hematopoietic organs *in vitro* (n = 7). (**H**) Scheme design. At stage for 5L2d (day 2 *Bombyx mori* Larvae of the fifth instar), the anterior pair of HPO-Wing complexes were dissected and cultured in a hanging drop of 10 μL culture system supplemented with JAK inhibitor AG490 (100 nmol/L) or vehicle DMSO for 72 h, and the released blood cells were collected. (**I**) Hemocytes were produced *in vitro*. Scale bars 200 μm. (**J**) Statistics of the number of hemocytes. The number of hemocytes in the DMSO group from the same individual was normalized. Data are shown as mean ± SEM. P-values of significance are calculated by Student’s T-test. The online version of this article includes the following source data and figure supplement(s) for figure 3: **Source data 1.** Source data for ***Figure 3B, Figure 3J, Figure 3—figure supplement 3C and Figure 3—figure supplement 3D.*** **Source date2.** Source data for ***Figure 3C.*** **Figure supplement 1.** The main cell type of STAT-positive hemocytes was the granulocyte.

We investigated the gene transcriptional levels of key members of the JAK/STAT pathway in hemocytes. The level of *STAT* gene mRNA in the CK group continued to increase with the development of the silkworm; the gene expression in the AM group was significantly up-regulated as compared with the CK group at 48 h to 96 h after modeling, but rapidly decreased to the level of CK group at 192 h (***Figure 3D***). The transcription level of *HOP*, a homologous gene of *Drosophila JAK*, showed a trend very similar to that of the *STAT* gene, and was significantly up-regulated in the AM group compared with the CK group at 48 h to 192 h after modeling (***Figure 3E***). We also studied the transcriptional levels of the negatively-regulated genes *SOCS2* and *SOCS6* downstream of the JAK/STAT pathway; although both genes were down-regulated in the AM group as compared with the CK group at the initial stage after modeling, they were up-regulated and significantly higher than the CK group in the later stages; just the temporal expression profile had a phase delay compared with *HOP* and *STAT* genes (***Figure 3F-G***).

In order to test the regulatory effect of JAK/STAT signaling pathway on silkworm hematopoiesis, highly congruent symmetrical HPO-wing complexes of 5L2d larvae, in which hematopoiesis was the most vigorous, were used for hanging drop culture on insect medium supplemented with JAK inhibitor AG490. The number of hemocytes in the culture medium after 72 h showed that AG490 significantly reduced the proliferation and release of hemocytes in HPO (***Figure 3I-J***). This supported the hypothesis that JAK/STAT signaling pathway promoted blood cell production in hematopoietic organs. This also showed that under the physiological environment of hyperproteinemia, the JAK/STAT signaling pathway in hemocytes was activated, and physiological processes such as regulation of blood cell proliferation are initiated.

### HPPC induced increasing phagocytic capacity and PCD levels of blood cells

In order to further explore the pathological mechanism by which HPPC affected blood cell homeostasis, the changes in the phagocytic ability of blood cells were investigated (***Figure 4—figure supplement 4***). The ink phagocytosis test showed that the blood cells swallowed ink particles and the number of ink spots in the AM group were more than those in the CK group, 48 h to 192 h after modeling (***Figure 4A***). The phagocytosis of fluorescent microspheres was further used to quantitatively detect the phagocytic ability of blood cells. On statistical analysis, we found that the ratio of phagocytic blood cells to the total blood cells in the AM group was about twice that of the CK group at the initial 48 h after modeling, then gradually decreased and returned to the control level at 192 h (***Figure 4D***). However, the phagocytic index (PI), which reflected the phagocytic ability of a single blood cell, was 44% to 75% higher in the AM group than in the CK group (***Figure 4C-E***). This demonstrated that HPPC improved the phagocytic ability of circulating blood cells.

**Figure 4.**
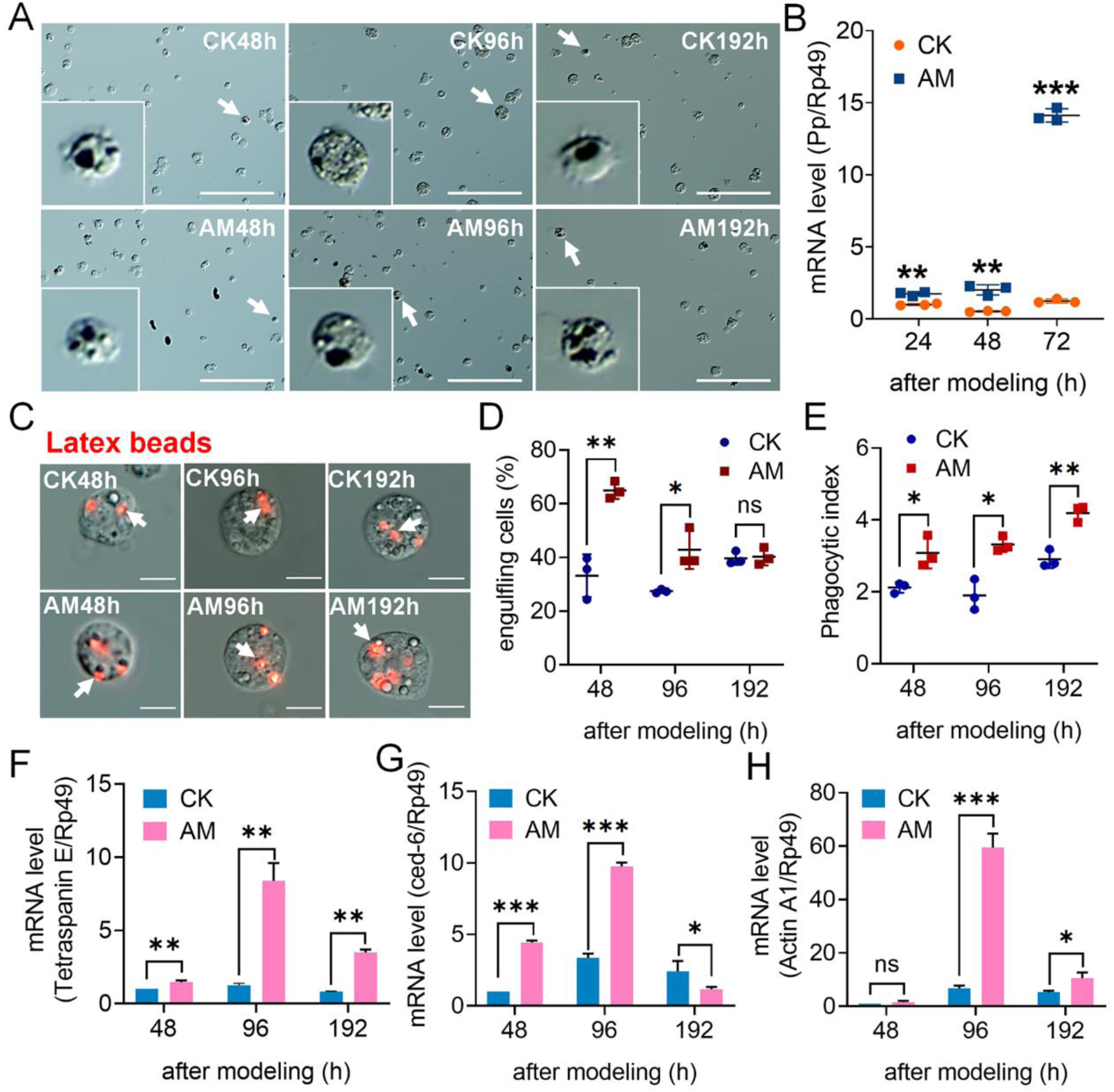
Effect of HPPC on the phagocytic ability of circulating hemocytes (blood cells). Ink or fluorescent microspheres were injected 48h, 96h or 192 h after modeling. The dose was 10 μL per silkworm pupae, and blood samples were collected 4 h later. (**A**) Ink phagocytosis test was used to qualitatively detect the phagocytic ability of blood cells (n = 3). Scale bars 100 μm. (**B**) The transcription level of the inflammatory factor Pp gene in hematopoietic organ-wing primordium complex (HPO-Wing) 24 h-72 h after modeling was analyzed by qPCR (n = 3). The internal reference gene was *Bombyx mori* Rp49. (**C-E**) The phagocytosis of fluorescent microspheres was used to quantitatively detect the phagocytic ability of hemocytes (n = 3). The number of fluorescent microspheres in 500-1000 hemocytes for each blood sample were counted by ImagePro software, and repeated 3 blood samples. (**C**) Characteristic diagram of hemocytes phagocytosing fluorescent microspheres. Scale bars 10 μm. (**D**) The percentage of phagocytes. (**E**) Phagocytosis Index (PI). PI = (number of particles phagocytosed by hemocytes / number of phagocytic hemocytes) × 100, reflecting the phagocytosis ability of a single hemocyte. (**F-H**) The mRNA level of phagocytosis-related genes in circulating hemocytes determined by qPCR (n = 3), and *Rp49* as an internal reference gene. Data are shown as mean ± SEM. ns p>0.05,*p<0.05, **p<0.01, and ***p<0.001 by Student’s T-test. The online version of this article includes the following source data and figure supplement(s) for figure 4: **Source data 1.** Source data for ***Figure 4D, Figure 4E and Figure 4—figure supplement 4A.*** **Figure supplement 1.** HPPC induced the increasing phagocytic capacity. **Figure supplement 2.** Hemocytes phagocytized (fat body) tissue fragments at 192 h after modeling.

We next investigated the mRNA level of genes *Tetraspanin E* (***Figure 4F***), *ced-6* (***Figure 4G***), and *Actin A1* (***Figure 4H***) as related to blood cell phagocytosis of silkworm; we found that the transcriptional level of *Actin A1* gene in AM group and CK were not significantly different at 48 h after modeling, and that the mRNA level of *ced-6* gene at 192 h after modeling was lower than that of the CK group. The transcriptional levels of the three genes in other time periods were significantly up-regulated compared with the CK group during the same period. This supported our results of enhanced phagocytic ability of blood cells in an HPPC physiological environment.

In order to explore the reasons for the enhanced phagocytic ability of blood cells in the AM group, we investigated the transcriptional level of the silkworm inflammatory factor *Pp* gene, which is specifically expressed in hematopoietic organs and affects cellular and humoral immunity. The results showed that during the entire study period, the level of *Pp* gene mRNA in the hematopoietic organ-wing primordium complex (HPO-Wing) of the AM group was significantly up-regulated compared with the CK group, and even reached 11.3 times that of the CK group at 72 h after modeling (***Figure 4B***). This showed that HPPC stimulated the immune response of hematopoietic organs.

We analyzed the mechanism by which HPPC led to changes in hemocyte homeostasis from the perspective of blood cell death. It was found that in the early and mid-term period (48 h to 96 h) after modeling, the autophagy signal in hemocytes was enhanced, with the level of autophagy increasingly being dominated by granulocytes; however, both decreased at 192 h. MDC staining (***Figure 5A***) marked the autophagy level of hemocytes. The MDC positivity rate of hemocytes in the AM group was significantly higher than that of CK before 96 h, but returned to the level of CK group at 192 h (***Figure 5B***). Regardless of the group (AM group or the CK group), the MDC-positive hemocytes in all investigation periods were mainly granular cells (***Figure 5C***). These were further stained with Lyso-Tracker Red (***Figure 5D***), and the results obtained showed a trend similar to MDC staining, that is, the autophagy-lysosome pathway (ALP) was activated significantly in the early and mid-term period (48 h and 96 h) after HPPC-induced modeling, but at 192 h, it returned to the level of the CK group (***Figure 5E***).

**Figure 5.**
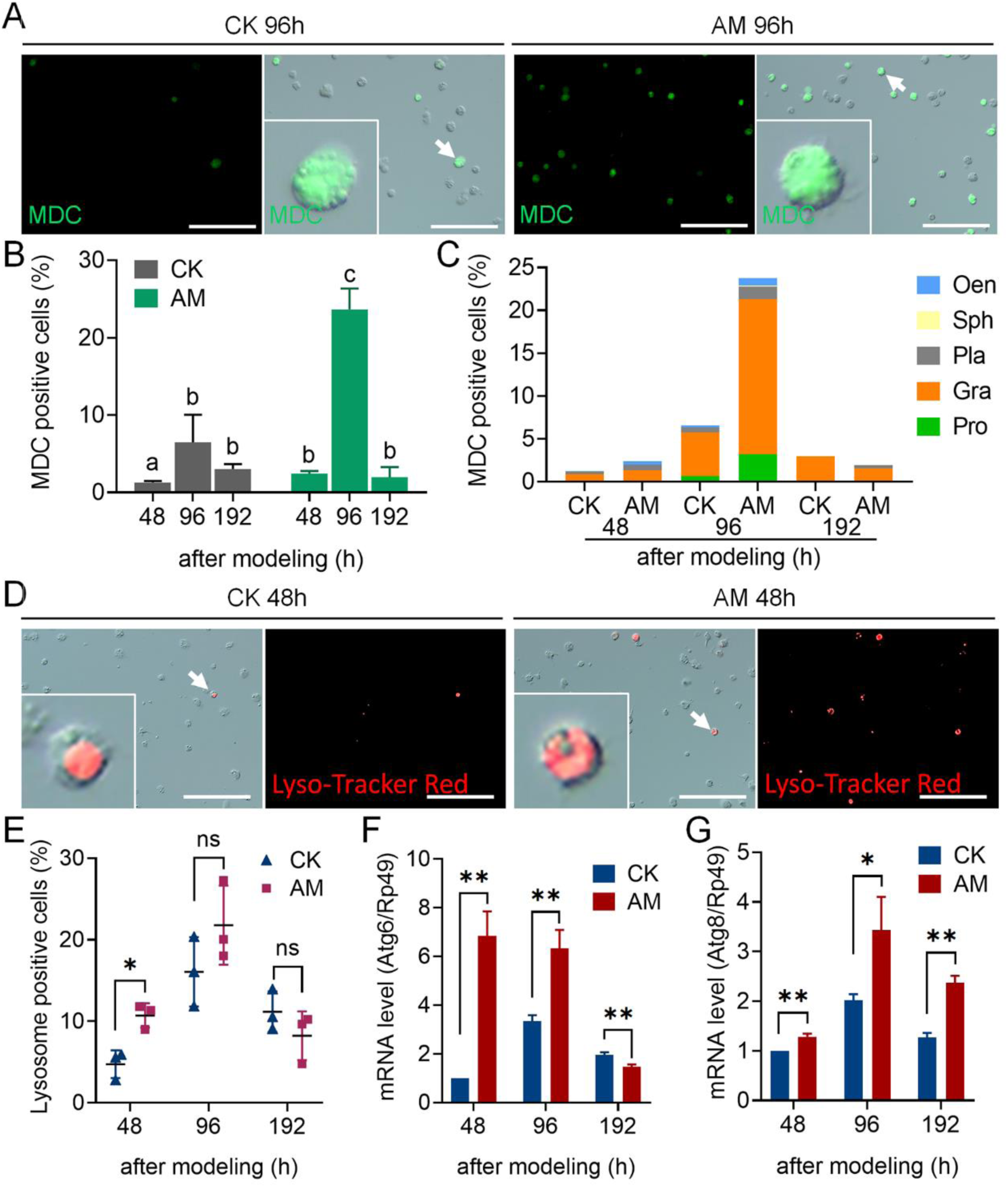
HPPC induced autophagy of circulating hemocytes (blood cells) in the early and mid-stage after modeling. MDC and Lyso-Tracker Red staining marked autophagy hemocytes in hemolymph at 48 h, 96 h and 192 h after modeling. Positive rate (%) = (fluorescent cell number / total cell number) × 100. Scale bars 100 μm. (**A**) Fluorescence image of MDC stained hemocytes 96 h after modeling (n = 3). (**B**) MDC staining positive rate of hemocytes. There were significant differences between groups with different letters (P≤0.05, n = 3). (**C**) MDC staining positive rate of different types of hemocytes. (**D**) Lyso-Tracker Red staining fluorescence image of hemocytes 48 h after modeling. Scale bars 100 μm. (**E**) Lyso-Tracker Red staining positive rate of hemocytes (n = 3). The transcription levels of autophagy genes in hemocytes (**F**) *Atg6* and (**G**) *Atg8* analyzed by qPCR (n = 3). The internal reference gene was *Bombyx mori* Rp49. Data are shown as mean ± SEM. ns p>0.05,*p<0.05, **p<0.01, and ***p<0.001 by Student’s T-test. The online version of this article includes the following source data and figure supplement(s) for figure 5: **Source data 1.** Source data for ***Figure 5B, Figure 5C and Figure 5E.***

We investigated the transcriptional level of marker genes in the autophagy signaling pathway. The expression of the upstream signal marker gene *Atg6* for autophagy initiation in AM silkworm hemocytes was significantly up-regulated as compared with the CK group, 48 h to 96 h after modeling, but at 192 h the expression completely recovered, and was even significantly down-regulated compared to the CK group (***Figure 5F***). However, the autophagy executive protein marker gene *Atg8* was up-regulated as compared with the CK group at 48 h to 192 h (***Figure 5G***).

TUNEL staining marks apoptotic hemocytes (***Figure 6A***). At 96 h after modeling, there was no significant difference between the positive cell rates of AM and CK, but both increased compared with levels at 48 h. At 192 h, the positivity rate of TUNEL in the CK group decreased from 9.12% at 96 h to 3.41%, and recovered to the level as at 48 h; while in the AM group it increased significantly from 9.64% at 96 h to 23.23% at 192 h, which was also significantly higher than that of the CK group (***Figure 6B***). On classifying the TUNEL-stained hemocytes, the apoptotic hemocytes were found to be mainly granulocytes in both the AM and CK groups (***Figure 6C***). We also studied the transcriptional level of *Dronc*, a marker gene of apoptosis initiation signal, and its level in hemocytes, and found that the level in the AM group was significantly higher than that in the CK group at 48 h to 96 h after modeling, but fell back to lower than the CK group level at 192 h (***Figure 6D***).

**Figure 6.**
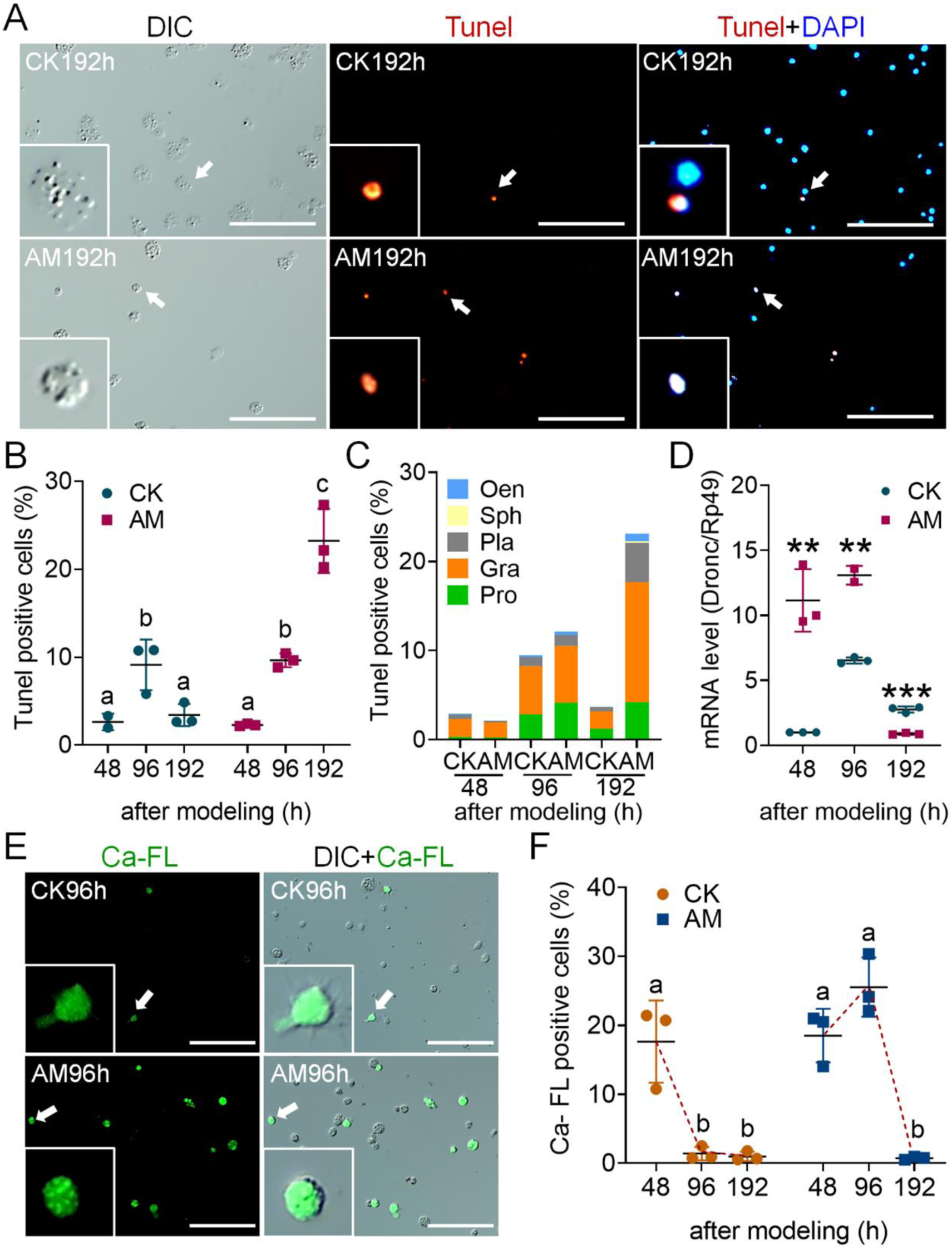
HPPC induced apoptosis of circulating hemocytes (blood cells) via the endoplasmic reticulum-calcium ion release signal pathway. Tunel staining and Ca-FL staining respectively marked the hemocytes with apoptosis and elevated cytoplasmic calcium level in the hemolymph 48 h, 96 h, and 192 h after modeling, and DAPI marked all hemocytes nuclei. Positive rate (%) = (fluorescent cell number / total cell number) × 100. Scale bars 100 μm. (**A**) 192 h Tunel stained fluorescence image after modeling. (**B**) Tunel staining positive rate of hemocytes. (**C**) Tunel staining positive rate of different types of hemocytes. (**D**) The transcription level of the apoptosis-initiating gene *Dronc* analyzed by qPCR, the internal reference gene was *Rp49* (n = 3). (**E**) Ca-FL staining fluorescence image of hemocytes 96 h after modeling. (**F**) The positive rate of hemocyte Ca-FL staining. In Figure 6B and Figure 6F, there were significant differences between groups with different letters (Data are shown as mean ± SEM, P≤0.05, n = 3). In Figure 6D, Data are shown as mean ± SEM. *p<0.05, **p<0.01 and ***p<0.001 by Student’s T-test. The online version of this article includes the following source data and figure supplement(s) for figure 6: **Source data 1.** Source data for ***Figure 6B, Figure 6C, Figure 6F, Figure 6—figure supplement 5B, Figure 6—figure supplement 5C and Figure 6—figure supplement 6B.*** **Figure supplement 1.** HPPC did not cause significant changes in circulating blood cell necrosis. **Figure supplement 2.** HPPC-induced apoptosis of blood cells did not depend on the mitochondrial-Caspase pathway.

We used Hoechst and PI co-staining to mark necrotic hemocytes (***Figure 6—figure supplement 5A***). At 48 h, 96 h, and 192 h after modeling, the co-positivity rates of hemocytes in the AM group were 3.42%, 2.97%, and 3.88%, respectively, compared with 3.37%, 2.96% and 3.13%, respectively, in the CK group (both at normal physiological levels) (***Figure 6—figure supplement 5B***). The co-positivity of hemocytes was further classified and investigated. At 48 h, the co-positive cells of the CK group and the AM group were mainly prohemocytes, granulocytes, plasmatocytes, and oenocytoid; at 96 h and 192 h after modeling, the co-positive cells in CK group were mainly prohemocytes, granulocytes, and plasmatocytes, while the AM group comprised mostly of the granulocytes (***Figure 6—figure supplement 5C***).

Fluo-3 AM fluorescent probe was further used to detect the level of apoptosis signal released by endoplasmic reticulum-calcium ion (***Figure 6E***). It was found that at 48 h after modeling, the positivity rates of Fluo-3 AM in hemocytes in the CK group and AM group were both high, reaching 17.64% and 18.52%, respectively. Subsequently, the positivity rates in the CK group quickly dropped to a low level, with only 1.39% and 0.99% at 96 h and 192 h, respectively; while in the AM group, the positivity rate rose rapidly to 25.55% at 96 h, but it also dropped to trace levels of 0.75% at 192 h (***Figure 6F***). Immunofluorescence of the apoptosis signal protein caspase-3 showed that the positivity rate of caspase-3 in hemocytes at 48 h to 192 h after modeling was very low; for the the AM group it was 0.60% to 1.58%, respectively, and for the CK group it was 0.77% to 3.19%, respectively (***Figure 6—figure supplement 6A-B***).

This showed that HPPC-induced apoptosis of blood cells did not depend on the mitochondrial-Caspase pathway, but mainly depended on the endoplasmic reticulum-calcium ion release apoptotic signaling pathway. Apoptosis occurred in the later stage of modeling after the autophagy signal and autophagy level fell back. The main apoptotic hemocytes included prohemocytes in addition to granulocytes.

On investigating the reasons for the increase in PCD levels in circulating hemocytes induced by HPPC, it was found that the blood ammonia level, that was closely related to HPPC, was always significantly higher in the AM group than in the CK group; levels were 28.29, 12.69, and 28.02 times that of the CK group at 96 h, 144 h, and 192 h, respectively, after modeling (***Figure 7—figure supplement 3B***). Using DCFH-DA staining to mark the ROS level of hemocytes, it is found that HPPC led to an increase in the rate of DCFH-DA positive hemocytes (***Figure 7A and Figure 7—figure supplement 3A***); the ROS level represented by the relative fluorescence intensity (ROS intensity) continued to rise after modeling, and was higher than that of the CK group (***Figure 7B***). After modeling, the H_2_O_2_ content (***Figure 7C***) and superoxide anion (ORF) level (***Figure 7D***) in the hemolymph of the AM group were several times that of the CK group, and the inhibitory ability of hydroxyl free radicals was also significantly lower than that of CK at 192 h (***Figure 7E***). The detection of related enzyme activities in hemolymph showed that at 48 h to 192 h after modeling, the activities of SOD (***Figure 7F***) and CAT (***Figure 7G***), and GSH content (***Figure 7H***) were significantly higher than those of the CK group. This indicated that hemolymphs increased the level of oxidative stress caused by the elimination of ROS, the decomposition of peroxides, and response to the adverse physiological milieu created by HPPC. This also showed that HPPC induced continuous and enhanced oxidative stress in hemocytes, with H_2_O_2_ being the main marker, which may be the reason for further increase in the PCD level of hemocytes.

**Figure 7.**
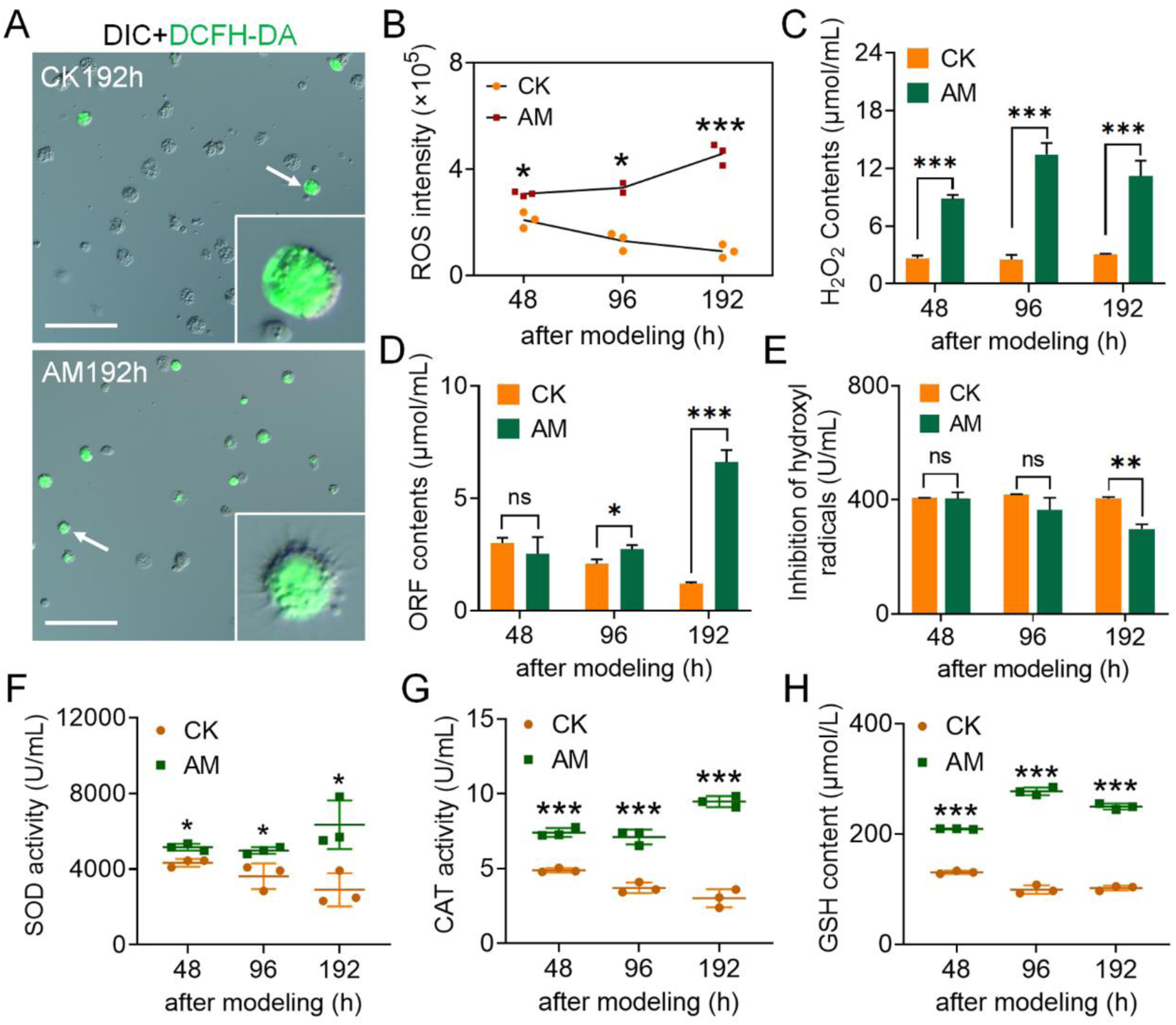
HPPC caused the oxidative stress of hemocytes to increase. DCFH-DA staining marked the ROS levels of hemocytes at 48 h, 96 h, and 192 h after modeling. Scale bars 100 μm. (**A**) DCFH-DA staining fluorescence image at 96 h after modeling. (**B**) The relative level of ROS in circulating hemocytes. Fluorescence intensity was calculated using Image Pro software, relative ROS intensity (× 10^5^) = fluorescence intensity / total hemocytes count. (**C-E**) Three main components of ROS in plasma. (**C**), H_2_O_2_ content; (**D**), superoxide anion (OFR) content; (**E**), hydroxyl radical (·OH) inhibition ability. (**F-H**) The activity of SOD and CAT, and GSH content in plasma. **A-H**, n = 3. Data are shown as mean ± SEM. ns p>0.05,*p<0.05, **p<0.01, and ***p<0.001 by Student’s T-test. The online version of this article includes the following source data and figure supplement(s) for figure 7: **Source data 1.** Source data for ***Figure 7B, Figure 7C, Figure 7D, Figure 7E, Figure 7F, Figure 7G, Figure 7H and Figure 7—figure supplement 3B.*** **Figure supplement 1.** HPPC increased the level of blood ammonia in hemolymph between 48 h and 192 h after modeling.

Next, we investigated the rescue effect of endocrine hormones on the homeostasis of circulating hemocytes caused by HPPC. By constructing a mild model mAM and injecting 20E rescue treatment 24 h after modeling, the PPC level of the rescue group was slightly lower than that of mAM group at 144 h after injection, but it was still significantly higher than in the CK group (***Figure 8A***). It is worth noting that the homeostasis of circulating hemocytes in the rescue group was significantly restored (***Figure 8B-C***).

**Figure 8.**
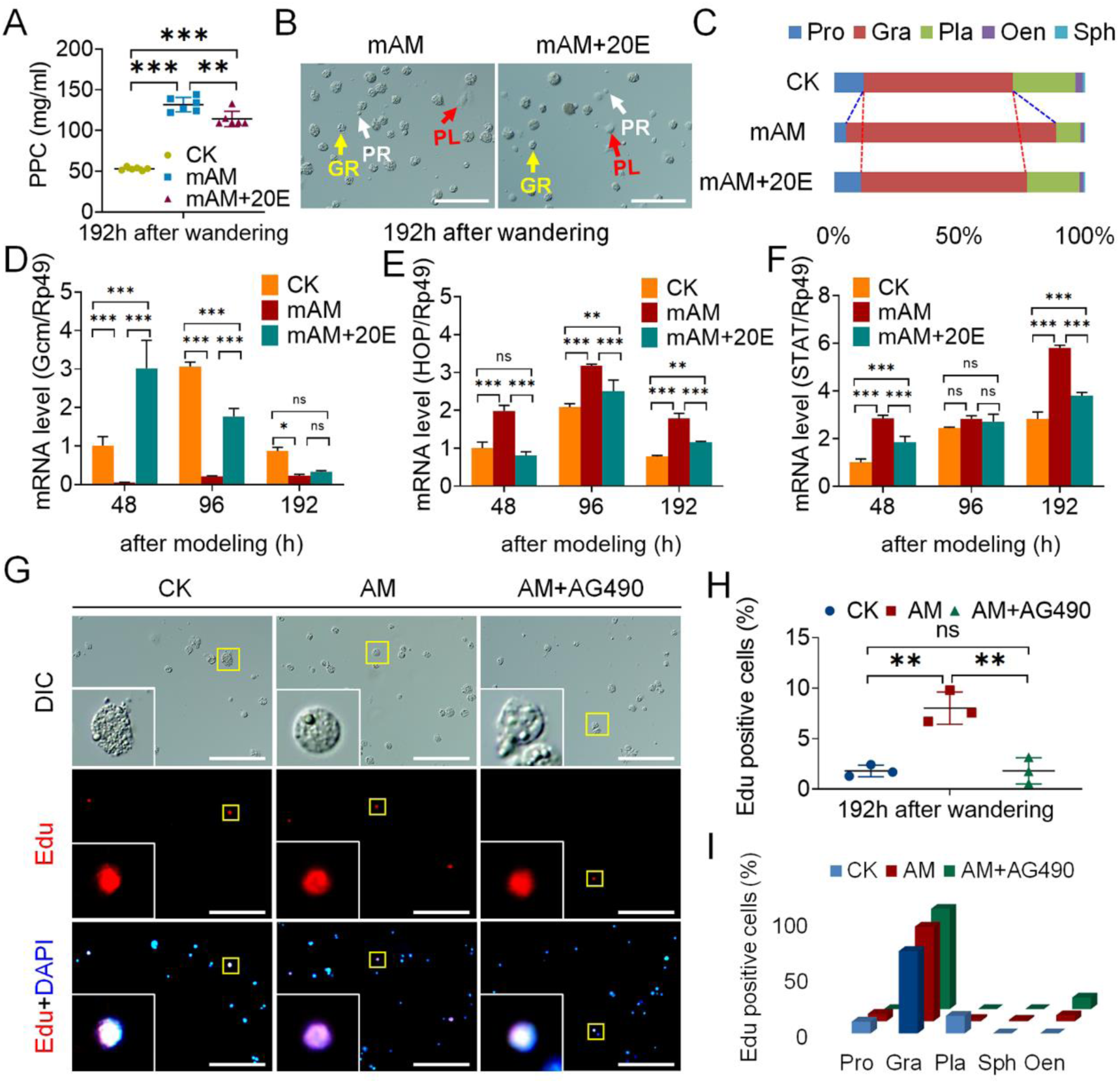
Endocrine hormone treatment of hyperproteinemia affected the proliferation and differentiation of hemocytes. **(A-C**) The percentage of hemocytes in the mAM group after 20E rescue. CK, control; mAM, mild model; mAM+20E, 20E was injected at 24 h after mAM modeling, and hemolymph samples were collected at 144 h after injection (corresponding to 192 h after AM group modeling). (**A**) PPC level (n = 5). (**B**) Circulating hemocytes. PR, Prohemocyte; PL, Plasmatocyte; GR, Granulocyte. Scale bars 100 μm. (**C**) Percentage composition of circulating hemocytes. (**D-F**) qPCR investigated the transcription level of related genes in the mAM group after 20E rescue (n = 3). The internal reference gene was *Bombyx mori* Rp49. mAM+20E, mAM group was injected with 20E at 0 h after modeling. (**G-I**) Edu staining was used to investigate the proliferation level of hemocytes after injection of JAK inhibitor A490 (n = 3). AM+AG490, the silkworm of AM group was injected with AG490 (50 μmol/10 μL) at 144 h after modeling, then the hemolymph samples were collected at 192 h. (**G**) Edu staining fluorescence image. (**H**) Edu-positive rate of blood cells. (**I**) Edu-positive rate of different types of hemocytes. Scale bars 100 μm. (A, D, E, F, H and I): Data are shown as mean ± SEM. ns p>0.05,*p<0.05, **p<0.01, and ***p<0.001 by Student’s T-test. The online version of this article includes the following source data and figure supplement(s) for figure 8: **Source data 1.** Source data for ***Figure 8A, Figure 8C, Figure 8H, Figure 8I, Figure 8—figure supplement 7D, Figure 8—figure supplement 7E, Figure 8—figure supplement 8 and Figure 8— figure supplement 9A.*** **Figure supplement 1.** The JAK / STAT signaling pathway activated by HPPC did not cause immune gene response. **Figure supplement 2.** Injection of AG490 in the late stage of hyperproteinemia enhanced the apoptosis of blood cells. **Figure supplement 3.** HPPC led to the synthesis and catabolism disorders of protein, sugar, lipid and other metabolites. **Figure supplement 4.** JAK/STAT pathway was activated in 4 patients with elevated PPC and accompanied by peripheral blood homeostasis imbalance.

Furthermore, 20E was supplemented at 0 h after the model of the silkworm in the mAM group. As a result, the hematopoietic factor Gcm in the hemocytes and the transcriptional levels of key genes in the JAK/STAT pathway showed changes similar to the CK group. In the hemocytes of the mAM group supplemented with 20E, the level of *Gcm* gene mRNA was significantly higher than that in the mAM group, and was notably higher than that in the CK group at 48 h after modeling (***Figure 8D***). The transcription levels of *HOP* and *STAT* genes that were significantly increased in the mAM group were significantly down-regulated after 20E supplementation (***Figure 8E-F***).

In order to further verify the effect of JAK/STAT pathway on the homeostasis of circulating hemocytes caused by HPPC, individuals in the AM group were injected with JAK inhibitor AG490 at 144 h after modeling, and circulating hemocytes were collected for investigation at 192 h. The results showed that the JAK/STAT pathway signal, marked with the *STAT* transcription level, is significantly weakened. Meanwhile, the Edu-positivity cell rate of proliferating cells, was significantly reduced from 8.03% in the AM group to 1.82% (***Figure 8G and Figure 8—figure supplement 7A***), and restored to the level of the CK group (1.80%) (***Figure 8H***). Classification of Edu-positive hemocytes revealed that the cell type that were decreased were mainly granulocytes (***Figure 8I***).

After injection of AG490, the occurrence of apoptosis of hemocytes in the AM group was significantly increased, and the number of surviving hemocytes was significantly reduced (***Figure 8—figure supplement 7D-E***). The results of the experiments with antimicrobial peptide genes showed that at 192 h after modeling, although the main antimicrobial peptide genes *moricin*, *Cecropin*, and *Defensin* in the hemocytes were significantly up-regulated in the AM group compared with the CK group, the mRNA levels were not significantly changed by AG490 injection (***Figure 8—figure supplement 7B***). This implied that the JAK/STAT signaling pathway activated in the later stage of HPPC was not the primary cause of immune gene response.

## Discussion

### Hyperproteinemia induces blood cell proliferation through the JAK/STAT signaling pathway

Studies have confirmed that hyperglycemia, hyperlipidemia, hypercholesterolemia, and other metabolic disorders affect the proliferation, differentiation, and death of blood cells (***Kraakman et al., 2017; Sarrazy et al., 2016; Nagareddy et al., 2013***), leading to changes in blood cell homeostasis and to hematopoietic disorders (***Barrett et al., 2019; Adler et al., 2014***). The effect of high cholesterol on blood cell homeostasis is related to the proliferation of multipotent progenitor cells (HSPC) and preferential differentiation into myeloid precursors (***Chistiakov et al., 2018***). ApoA-I binding protein-mediated cholesterol efflux activates endothelial Srebp2 and promotes the expansion and mobilization of HSPCs by regulating Notch pathway genes (***Gu et al., 2019***). The proliferation and migration of myeloid progenitor cells is induced by hyperglycemia, which is related to the effect of S100A8/S100A9 produced in neutrophils and induced by monocytes on the receptors for advanced glycation end products on myeloid progenitor cells (***Nagareddy et al., 2013***). Hyperlipidemic serum promotes the production of increased IL-6 by immunocytes called dendritic cells (DCs), thereby enhancing the production of IL-17A, leading to an increase in granulocytes (***Bagchi et al., 2017***). AMPK activated by M-CSF accelerates atherosclerosis by enhancing the differentiation of monocytes into macrophages (***Zhang et al., 2017***). It can be seen that although the pathological mechanisms of derangement in blood cell homeostasis caused by different metabolic disorders are different, they are all related to the conservative hematopoietic transcriptional factors and the dysregulation of gene expression in signaling pathways, such as M-CSF, AMPK, and Notch (***Gu et al., 2019; Zhang et al., 2017***).

The hematopoietic function of the silkworm is regulated by a series of conserved hematopoietic factors and signaling pathways, including *Ush*, *Lozenge*, and *Gcm* (Gene ID: 101742696), which is the homologous gene of *DmGcm* of *Drosophila*. *Ush* and *Lozenge* are involved in regulating the production of oenocytoids (***Xu et al., 2015***), and the *DmGcm* is found to regulate the differentiation and formation of plasma cells in *Drosophila* (***Bataillé et al., 2005; Lebestky et al., 2000***). Down-regulation of *DmGcm* gene expression will enhance the phenotype caused by overexpression of the JAK/STAT pathway (***Trébuchet et al., 2019; Bazzi et al., 2018***). In mammals and fruit flies, highly conserved signaling pathways such as JAK/STAT are necessary to maintain normal blood cell homeostasis (***Koranteng et al., 2020; Raivola et al., 2019; Hao and Jin, 2017***). When there is an imbalance in blood cell homeostasis due to leukemia or diabetes, dysregulation of the cellular JAK/STAT pathway has been observed (***Huang et al., 2018; Dodington et al., 2018***). Using a cultured cell line BmN, they found that JAK/STAT may be involved in the regulation of the silkworm cell cycle (***Hu et al., 2015***). Furthermore, it has an interaction with the epidermal growth factor receptor pathway in silkworm larvae (***Abbas et al., 2018***).

This study found that HPPC caused changes in the basal metabolism of hemolymph in *Bombyx mori*, induced a significant increase in the activity level of the endocrine hormone bombyxin, which regulated glucose metabolism, and led to the synthesis and catabolism disorders of protein, sugar, lipid and other metabolites (***Figure 8—figure supplement 8A-I***). HPPC severely and adversely affected the homeostasis of blood cells, leading to a decrease in the number of prohemocytes with multidirectional differentiation potential in the circulating hemolymph, and a significant increase in granulocytes that exerted phagocytic ability; but the other phagocytic blood plasmatocytes were significantly reduced. This showed that hematopoietic disorder was an important reason for the imbalance of blood cell ratio caused by primary hyperproteinemia. Further mechanistic studies have found that the effect of HPPC on hematocyte proliferation and differentiation was closely related to the increase in JAK/STAT signal levels and the significant down-regulation of *Gcm* transcriptional levels. After supplementation with 20E hormone therapy, *Gcm* expression level of hematopoietic factor was restored in blood cells with down-regulation of JAK/STAT signal level; moreover, the imbalance in circulating blood cell composition also improved. This implied that endocrine hormones reversed the activation of related molecular pathways and provided rescue from the effects of HPPC on the homeostasis of circulating hematocytes. This, therefore, suggested that HPPC activated a signaling pathway that affected blood cell homeostasis, by a mechanism which was different from the effects of high blood sugar, high blood fat, and high cholesterol.

Clinical studies on multiple myeloma that is often secondary to hyperproteinemia have found that the JAK/STAT3 signaling pathway, which is abnormally activated by IL-6, played an important role in promoting the survival, proliferation, and drug resistance of myeloma cells (***Chong et al., 2019; Ren et al., 2019***), but there is still a lack of in-depth research on the influence or mechanism of action on peripheral blood cells. In this study, a survey of four clinical samples with elevated PPC, including patients with multiple myeloma, found that compared with normal PPC samples, JAK/STAT pathway genes in blood cells were activated to varying degrees, and the expression level of transcription factor *Gcm* appeared to be significantly reduced, with an imbalance in peripheral blood homeostasis (***Figure 8—figure supplement 9A-B***). This observation further supported the results that HPPC in silkworms activated the JAK/STAT signaling pathway of circulating blood cells.

**Figure 9.**
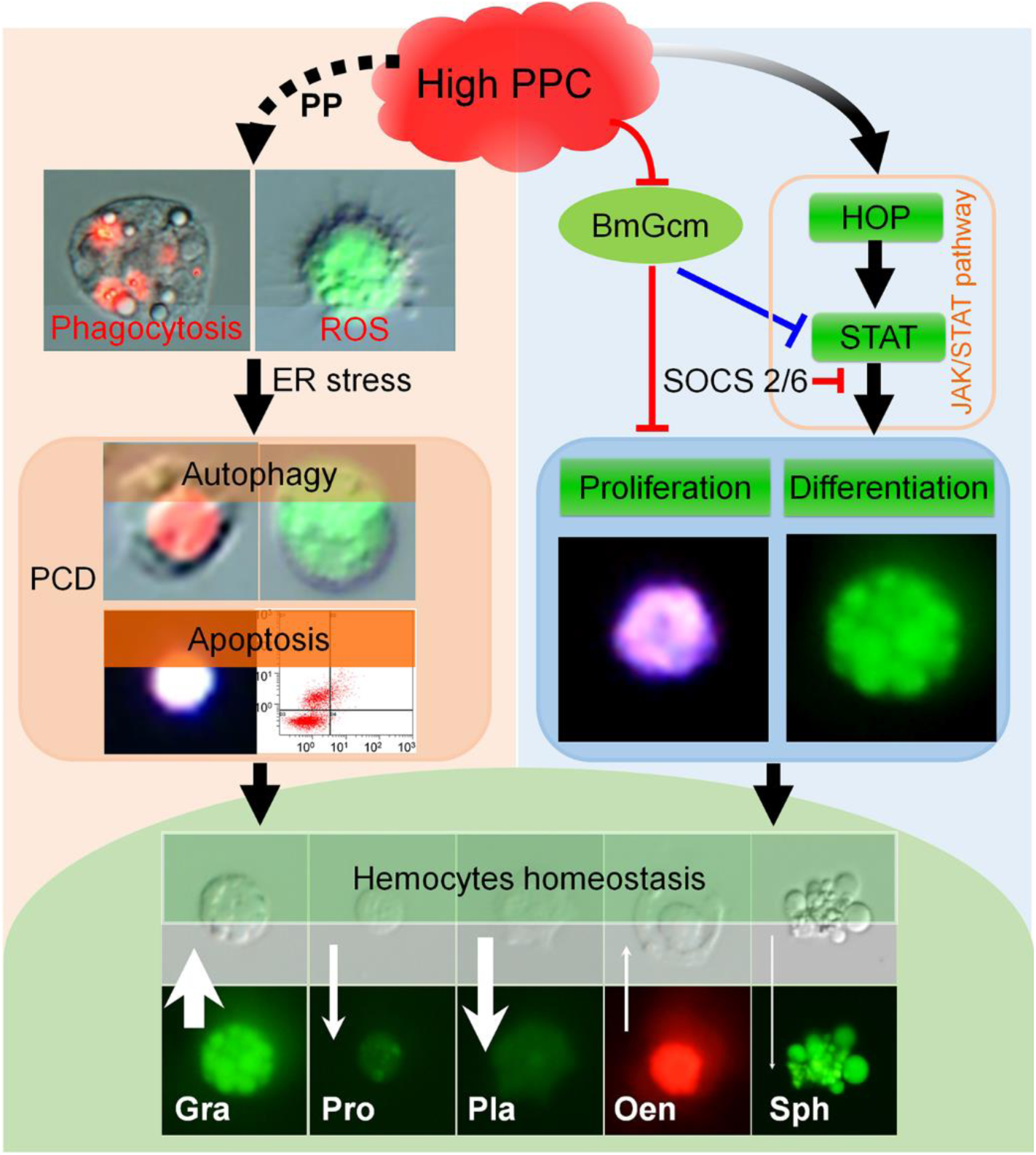
Summary of HPPC affecting blood cell homeostasis. HPPC causes the phagocytic ability of hematocytes and the level of ROS to increase, and then induces the PCD process of hematocytes via the endoplasmic reticulum-calcium ion release signal pathway. HPPC activates the JAK/STAT signaling pathway and induces the proliferation of hematocytes dominated by granulocytes; the significant down-regulation of the *Gcm* gene further aggravates the process, which together affects the homeostasis of circulating hematocytes, resulting in an increase in the percentage content of granulocytes and oenocytoids, while decrease in that of prohemocytes, plasmatocytes and spherulocyte. The black solid line arrow indicates the positive correlation induction effect, and the black dotted line arrow indicates the correlation promotion effect; the red and blue stop lines indicate the inhibition (reverse regulation) effect of the results of this article and the results of the cited literature respectively. The white arrow indicates the change in the number of hematocytes in the hemolymph induced by HPPC, and the length of the arrow indicates the changed maximum ratio of density between two groups of the model and the control, and that of Gra, Pro, Pla, Oen, Sph were 2.38, 3.99, 9.82, 2.71 and 18.77 times, respectively. The width of the arrow indicates the absolute value of the maximum difference between the model and the control in the percentage of various types of hematocytes (maximum AM-CK), and that of Gra, Pro, Pla, Oen and Sph were 27.16%, 10.99%, 21.28%, 1.90% and 0.79%. The upward arrow indicates that AM is higher than CK, and the downward arrow indicates that AM is lower than CK.

### Hyperproteinemia induces programmed death of blood cells through the endoplasmic reticulum-calcium ion release signaling pathway

The phagocytic behavior of silkworm hemocytes mainly depends on granulocytes and plasmatocytes (***Ling et al., 2005***), and is accompanied by up-regulation of the expression of phagocytosis-related genes *ced-6*, *ActinA1*, and *Tetraspanin E* (***Li et al., 2017; Ishii et al., 2010***). The paralytic peptide Pp, which is specifically expressed in the hematopoietic organ-wing imaginal disc complex, is an inflammatory cytokine that mainly stimulates cellular and humoral immune responses (***Ishii et al., 2013; Ishii et al., 2010***). In this paper, HPPC caused a significant up-regulation of the *Pp* gene transcriptional level of the silkworm; meanwhile the phagocytic ability of circulating hemocytes enhanced significantly with the granulocytes rising rapidly. We, thus, demonstrated that the silkworm HPPC model also exhibited an increase in macrophages and enhanced phagocytosis associated with inflammation that occur in mammalian hyperglycemia and hypercholesterolemia (***Jaitin et al., 2019; Recio et al., 2018; Tall and Yvan-Charvet, 2015***).

Clinical studies have shown that blood cell PCD is a common response of the body to hyperglycemia, atherosclerosis, hyperhomocysteinemia, and other metabolic diseases (***Simion et al., 2020; Vion et al., 2017; Xi et al., 2016***); the molecular mechanisms induced include Caspase-dependent mitochondrial pathways and endoplasmic reticulum stress, and death receptor-mediated apoptosis pathways (***Mattisson et al., 2017; Xi et al., 2016***). In addition, members of ROS, such as hydrogen peroxide, hydroxyl radicals, and superoxide anions, resist the adverse external environment, and promote the bactericidal activity of phagocytes (***Laforge et al., 2020; West et al., 2011***); however, excessive production of ROS will cause continuous inflammation and promote blood cell apoptosis (***Silwal et al., 2020; Cui et al., 2018***).

The PCD pathway of silkworm hematocytes mainly depends on the exogenous apoptosis pathway mediated by death receptors expressed in the plasma membrane and the intrinsic apoptosis pathway controlled by mitochondria (***Galluzzi et al., 2018; Fuchs and Steller, 2015***). In this paper, along with the development of HPPC, autophagy and apoptosis of hematocytes were observed in the silkworm model, while cell necrosis was always at a low level (***Figure 6—figure supplement 5A-B***). At the same time, it was found that HPPC caused severe oxidative stress in hematocytes, and the level of the ROS, hydrogen peroxide (the main marker), increased. Mechanistic studies have found that HPPC caused an increase in the level of calcium ions in hematocytes, which was typically time-dependent with the occurrence of blood cell apoptosis. This showed that PCD of hematocytes induced by the apoptotic signaling pathway of endoplasmic reticulum-calcium ion release was an active response.

## Conclusion

HPPC caused the phagocytic ability of blood cells and the level of ROS to increase, and then induced the process of PCD of hematocytes via the endoplasmic reticulum-calcium ion release signaling pathway. On the flip side, HPPC activated the JAK/STAT signaling pathway of hematocytes to induce the proliferation of hematocytes, mainly granulocytes, and inhibited the PCD level of cells (***Figure 9***). Supplementation with 20E improved hematopoietic function and restored the homeostasis of circulating hematocytes by modulating the JAK/STAT signaling pathway.

## Materials and Methods

### Preparation of animals

The animal model of HPPC (AM) used by our group (***Chen et al., 2018***), and the processing time axis are shown in ***Figure 1—figure supplement 1A***. The larvae of silkworm species *Dazao* were fed with fresh mulberry leaves. The rearing environment of whole generation was: temperature, 25°C; relative humidity 70–80%, and 12 h-light followed 12 h-dark cycle. In the wandering stage, mature female larvae were selected, and low melting point wax (55°C–60℃) was used to seal the silk holes in the front of the head to prevent spinning. The model of *Bombyx mori* with significantly increased PPC was constructed by releasing nontoxic silk protein into hemolymph during the larval-pupal metamorphosis. In addition, 24 h after the start of silking, mature larvae were selected to plug the silking hole to prepare another mild model silkworm (mAM) with high PPC level that was non-lethal. The PPC level after modeling was about 60% of the AM group, and the same batch of naturally spinning silkworms were used as controls (CK). The animals of the mAM group were injected with 20E (20-hydroxyecdysone) (10 μL, 0.4 μg/μL per individual) (Sigma-Aldrich, Santa Clara, California, USA) at 0 h or 24 h after modeling, and the animals of AM group were injected with AG490 (JAK inhibitor) (10 μL, 50 μM per individual) (S1509, Beyotime, Jiangsu, China) at 144 h after modeling. After disinfection with 75% ethanol at the scheduled time, the epidermis was punctured, and the hemolymph was collected with an eppendorf tube in an ice bath (added with 10 μL saturated phenylthiourea solution), and equal volumes of hemolymph from 3–6 individuals were mixed to form a hemolymph sample for testing.

### Determination of phagocytic ability of blood cells

The phagocytic ability of blood cells (hemocytes) was qualitatively detected by using neutral carbon ink diluted 10 times (v/v) with PBS. The phagocytic capacity of hemocytes was quantitatively detected by the method used previously (***Ishii et al., 2010***). Fluorescent latex beads [FluoSpheres™ Carboxylate-Modified Microspheres, 0.1 µm, red fluorescent (580/605 nm, 2% solids) (F8801, Thermo Fisher Scientific, Waltham, Massachusetts, USA)] diluted thrice (v/v) with PBS were injected into each individual, and hemolymph sample (from six individuals) were collected 4 h later. The hemocytes were climbed and fixed similarly using TUNEL after the hemolymph samples were diluted. Lastly, the hemocytes were observed under fluorescence microscope (Olympus BX51, Olympus, Tokyo, Japan), and the number of fluorescent latex beads in hemocytes was counted by Image-Pro Plus 6.0 software (Media Cybernetics, Rockville, Maryland, USA).

### Blood cell classification

Using the method previously employed (***Ling et al., 2003***), the hemocytes of *Bombyx mori* were classified by acridine orange-propidium iodide (AO-PI) staining. To 40 μL of hemolymph we added 10 μL AO (10 μg/mL) (A6014, Sigma-Aldrich) and 10 μL PI (2 μg/mL) (P4170, Sigma-Aldrich) working solution, mixed well, and allowed the mixture to stand for 2 min. Red fluorescence (535/615 nm) and green fluorescence (488/515 nm) were observed under the fluorescence microscope.

### *In vitro* culture of hematopoietic organs

We referred to the literature for the method of *in vitro* culture (***Liu et al., 2014***). We used 75% ethanol to anesthetize the larvae on the second day of the 5th instar or 24 hours after modeling, and then took out the complete two front-end complex wing-hemopoietic organ (HPO), followed by washing with PBS and Grace’s Insect Medium. The single HPO was cultured in 10 μL medium at 26℃ in a hanging drop. The culture medium was Grace’s insect medium (11605094, Thermo Fisher Scientific) supplemented with 10% (v/v) silkworm serum, 1% (v/v) phenylthiourea saturated solution, and appropriate amount of antibiotics.

### Blood cell (hemocyte) staining

#### Edu staining to detect DNA replication in blood cells

Each individual was injected with 10 μL Edu: PBS (3 mg/mL). After 8 h, the hemolymph was collected and diluted 5–10 times (v/v) with HbSS. Furthermore, 60 μL of diluted hemolymph was taken for hematocyte climbing test (***Qiu et al., 2019***); we stained the hemocytes using Click-iT™ EdU Cell Proliferation Kit for Imaging, Alexa Fluor™ 594 dye (C10339, Invitrogen, Carlsbad, CA, USA). The cell nucleus was counter-stained with DAPI (C1006, Beyotime); red (590/615 nm) and blue fluorescence (364/454 nm) were observed microscopically. Control group was injected with 10 μL DMSO: PBS (1:2).

#### Reactive oxygen species (ROS) staining

Referring to the previous literature (***Li et al., 2017***), we took 100 μL of hemolymph diluted 5–10 times in normal saline, and climbed the film for 5 min. According to the instructions of the kit (S0033S, Beyotime), we added 1,000 μL ROS staining working solution (10 μM), incubated at 25°C for 30 min in the dark, washed with saline three times in the dark, and observed the green fluorescence (488/525 nm) under the fluorescence microscope.

#### Detection of blood cell cycle phase distribution

For this we used propiridine iodide-flow cytometry (PI-FCM) (***Qiu et al., 2019***). First, 600 μL cold HbSS (C0218, Beyotime) was added to 600 μL hemolymph sample to collect the hemocytes after centrifugation at 3,000 rpm at 4℃ for 10 min. Then, after HbSS washing and centrifugation twice, 1 mL of 70% cold ethanol was gently added and the hemocytes were fixed at 4℃ for 12 h. Finally, the hemocytes were washed twice with HbSS, mixed with 400 μL of PI working solution (C1052, Beyotime) and incubated at 37℃ for 30 min in dark. Cell cycle analysis was done using FC500 FCM (Beckman Coulter, Brea, California, USA).

#### Detection of blood cell autophagy

We used two methods for this (***Li et al., 2017***): Monodansylcadaverine (MDC) (30432, Sigma-Aldrich) and Lyso-Tracker Red (C1046, Beyotime). For MDC staining, after the hemolymph samples were diluted, the hemocytes were first climbed and fixed in a way similar to detection of apoptosis by TUNEL. After washing with PBS for three times, we added 1 mL of MDC working solution (50 μM) and incubated in the dark for 30 min. Green fluorescence (338/500 nm) was observed under the microscope. For lysosome staining, the hemocytes were first climbed for 5 min, then washed twice with PBS; we added 1,000 μL Lyso-Tracker Red working solution (0.1 μM) to cover the hemocytes, and incubated at 25℃ for 30 min. After washing three times with PBS, red fluorescence (577/590 nm) was observed under the microscope.

#### Detection of blood cell apoptosis

First, 60 μL of hemolymph samples were diluted 5–10 times (v/v) with HbSS. After 5 min of climbing, the hemocytes were fixed with 4% paraformaldehyde for 15 min; we used three methods to detect apoptosis. For detection using TUNEL (***Li et al., 2017***), hemocytes were incubated with PBS containing 0.5% Triton X-100 (T8787, Sigma-Aldrich) for 20 min at 25°C. This was followed by detection using the In Situ Cell Death Detection Kit, TMR red (12156792910, Roche, Basel, Switzerland) program, in which samples were incubated with 60 μL of reaction working solution in the dark at 37°C for 60 min. After DAPI counterstaining of the nucleus for 20 min, we observed red (540/580 nm) and blue fluorescence under the microscope. Calcium ion fluorescent probe was used to detect calcium ion concentration in the cytoplasm (***Li et al., 2017***). We added 1,000 μL Fluo-3 AM (S1056, Beyotime) staining solution (5 μM), incubated for 30 min in the dark at 25°C and observed the green fluorescence (488/530 nm) after staining. Apoptosis was also detected by FCM (***Qiu et al., 2019***); hemocytes were collected according to the method described above for PI-FCM, and the percentage of apoptotic blood cells was detected by Annexin V, FITC Apoptosis Detection Kit (AD10, DOJINDO, Tokyo, Japan) and FC500.

#### Immunofluorescence staining

After climbing, fixing, and permeabilizing blood cells in the same way as done in TUNEL, we added the primary antibody Cleaved Caspase-3 (Asp175) Rabbit Ab (9661s, Cell Signaling Technology, Boston, Massachusetts, USA) or purified anti-STAT (peptide sequence: LRKIKRAEKKGTESC) (1:200), and incubated for 12 h at 4℃. After washing with PBS, it was incubated with the secondary antibody Goat Anti-Rabbit IgG (H+L) FITC Conjugated (GAR001, MUTI SCIENCES, China) (1:200), and finally the nucleus was counter stained with DAPI; the green fluorescence (495/515 nm) of the target protein and blue fluorescence (364/454 nm) of the nucleus were observed under the microscope.

### Western blotting

Blood cells were collected from 5 mL hemolymph samples at 5,000 rpm at 4℃ for 10 min. Then, proteins were extracted by 100 μL RIPA Lysis Buffer (P0013B, Beyotime) containing 1% Phenylmethanesulfonyl fluoride (PMSF, Beyotime) and 1% phosphatase inhibitors (PhosSTOP, Sigma-Aldrich). Using TGX Stain-Free FastCast Acrylamide Kit (1610183, Bio-Rad, Hercules, California, USA), the protein samples were separated by 10% sodium dodecyl sulfate (SDS)–polyacrylamide gel electrophoresis, and transferred to polyvinylidene difluoride (PVDF) membrane using a semi-dry transfer film. The membrane was blocked with western blocking solution (P0023B, Beyotime) for 2 h at 25°C, followed by incubated with the purified anti-STAT (1:200) antibody and anti-Tubulin (ab52901, Abcam, Cambridge Science Park, England) (1:5000) for 12 h at 4°C. After washing three times with TBS containing 0.05% Tween 20 (TBST; pH 7.5), the membrane was incubated with HRP-labeled anti-rabbit IgG (Bioworld Technology, Minnesota, USA) (1:5000) for 2 h at 4°C. After washing three times with TBST, according to 1:1 by adding appropriate amount of ECL (Bio-Rad) color solution in the dark conditions and photographed using an EZ-ECL chemiluminometer Detection Kit for HRP (Biological Industries, Israel) after 2 min at room temperature. The resulting pictures were processed and analyzed using Image Lab software.

### Gene expression analysis

Total RNA was extracted using TRIzol^®^ Reagent, Invitrogen™ (15596018, Life Technologies, Carlsbad, California, USA) kit, and cDNA was obtained by reverse transcription. ABI StepOnePlus™ real-time PCR system (Ambion, Foster City, CA, USA) was used for qRT-PCR in a 20 μL reaction system. Gene primers are listed in ***Supplementary file 1***, and *Bombyx mori* Rp49 was used as the reference gene.

### Gene silencing

Small interfering RNAs (siRNAs) to *Gcm* (Gene ID: 101742696) were designed and synthesized by Genepharma (Shanghai, China); the sequence of primers (siRNA-Gcm 598 and siRNA-Gcm 1017) are shown in ***Supplementary file 1***. CK and mAM were injected with 10 μg of siRNA (siRNA-Gcm598: siRNA-Gcm 1017=1:1) at 96 h after modeling, and the negative control was injected with the same amount of negative control siRNA (si-NC). Hemolymph samples were collected 48 h after injection, and the interference efficiency was detected by qPCR.

### Biochemical assays

Hemolymph samples were collected from 48 h through 192 h after modeling, and the manual method of the kit (BC4385, BC3595, BC1290, Solarbio; A018, Nanjing Jiancheng, Jiangsu, China) was used to detect the levels of blood ammonia, hydrogen peroxide, superoxide anion, and hydroxyl free radicals. The enzyme activities of catalase (CAT) and superoxide dismutase (SOD), and glutathione S-transferase (GST) content were measured using the kit (A007; A001; A006, Nanjingjiancheng, China). The absorbance of all samples were measured using a microplate spectrophotometer (Eon, BioTek, Winooski, Vermont, USA).

### Data analysis

Image-Pro Plus 6.0 software and GraphPad Prism 8 software (GraphPad, San Diego, California, USA) were used for image processing and data processing, respectively. Holm-Sidak method for multiple *t*-tests (one per row) analysis was done.

## Acknowledgements

We thank Dr. Weimin Yin and Dr. Jianfen Qiu from Soochow University for their constructive suggestions.

## Additional information

### Funding

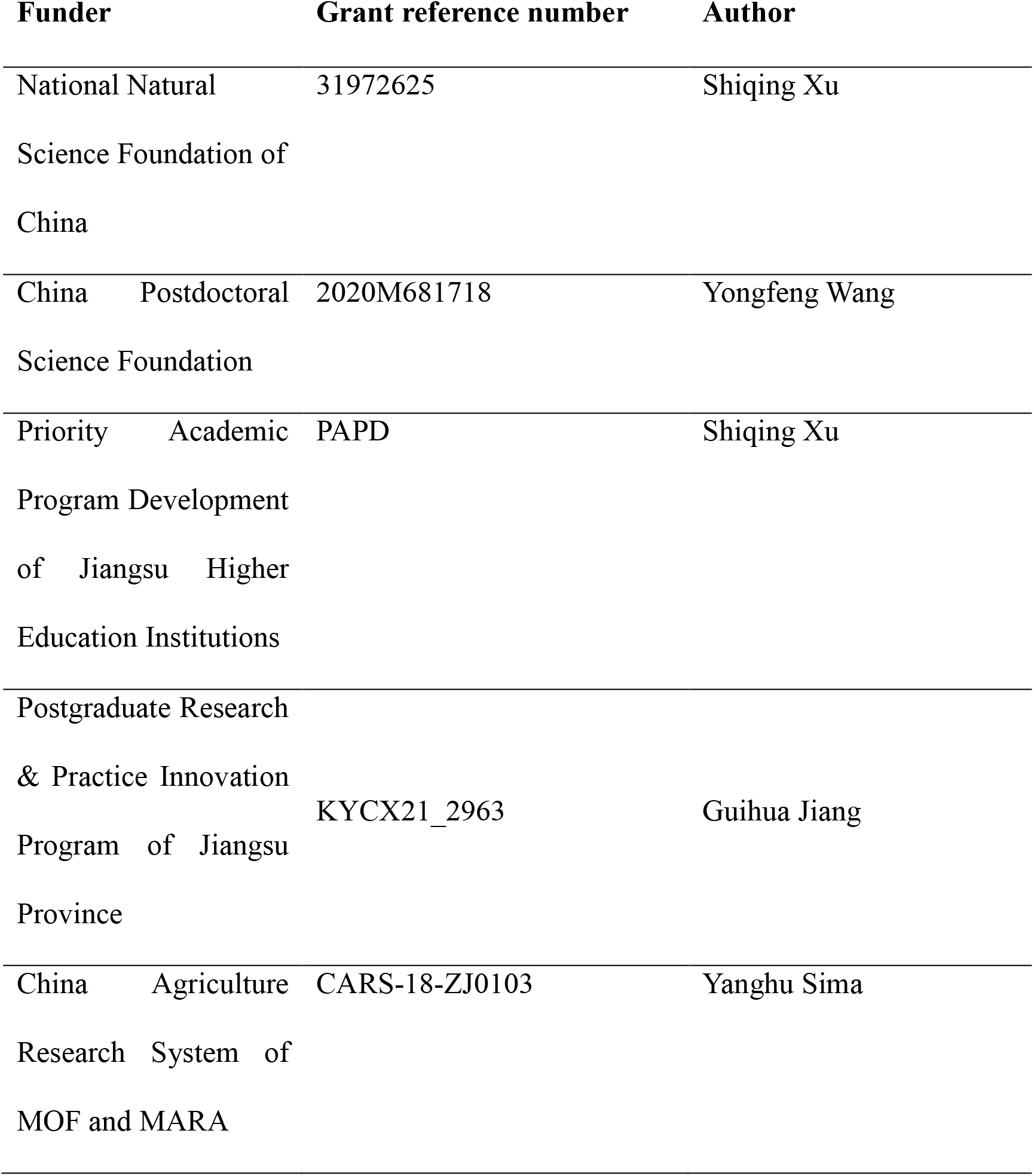
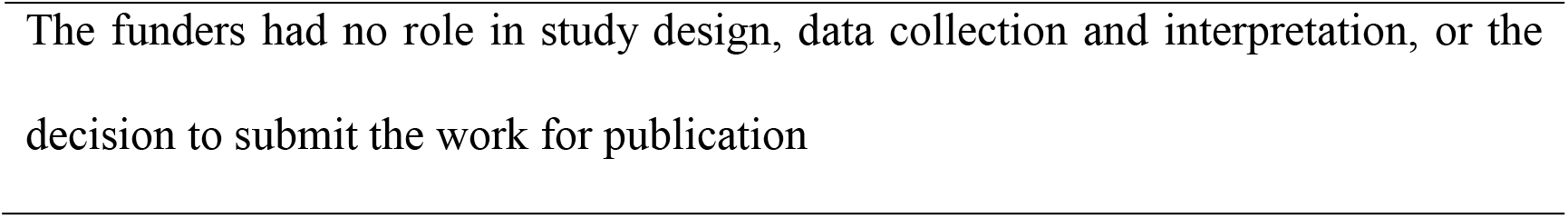

### Author Contributions

Guang Wang, Conceptualization, Data curation, Formal analysis, Investigation, Methodology, Project administration, Validation, Visualization, Writing – original draft, Writing – review and editing; Yongfeng Wang, Conceptualization, Data curation, Funding acquisition, Methodology, Writing – original draft; Jianglan Li, Investigation; Ruji Peng, Investigation; Xinyin Liang, Investigation; Xuedong Chen, Methodology; Guihua Jiang, Funding acquisition, Investigation; Jinfang Shi, Resources, Validation; Yanghu Sima, Funding acquisition, Resources, Supervision; Shiqing Xu, Conceptualization, Data curation, Formal analysis, Funding acquisition, Methodology, Project administration, Resources, Supervision, Validation, Writing – original draft, Writing – review and editing.

### Ethics

Animal experimentation: All silkworm rearing was performed under standard conditions, and all animal experiments were performed in strict accordance with the recommendations in the Guide for the Care and Use of Laboratory Animals of the National Institutes of Biological Sciences. All of the animals were handled according to the guidelines of the Chinese law regulating the usage of experimental animals and the protocols (#SUDA20210802H02 ) approved by the Committee on the Ethics of Animal Experiments of the Soochow University.

## Additional files

### Supplementary files

• Supplementary file 1. Primers used in this part.
• Supplemental Materials
• Transparent reporting form

### Data availability

All data generated or analyzed during this study are included in the manuscript and supporting files.

## Figure Supplements

**Figure supplement 1.**
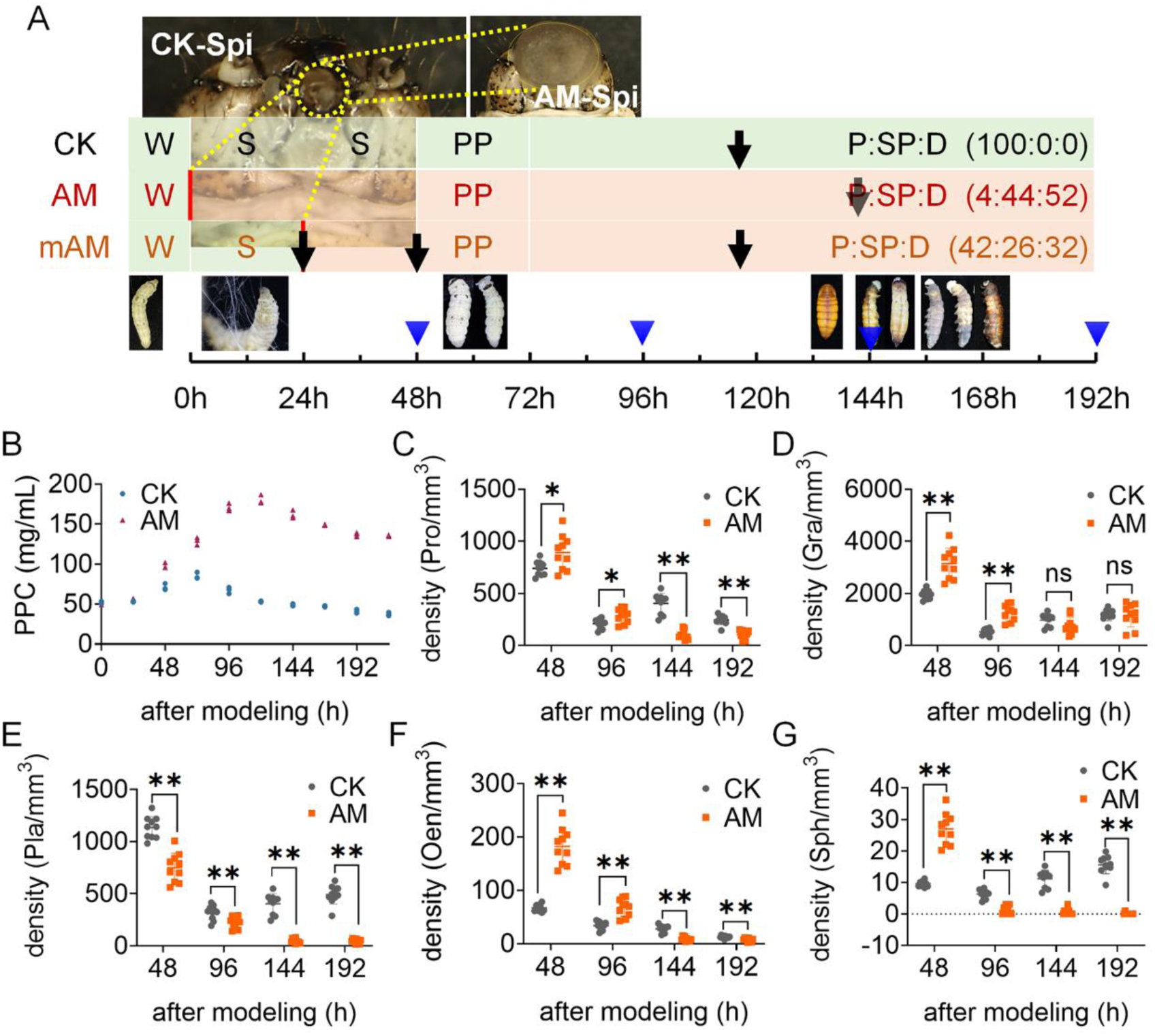
The effect of hyperproteinemia on the density of different types of hemocytes. (**A**) Timeline. The silkworm (Dazao) was reared to mature larvae (wanderding, W) in a standardized manner. Spi, spinnert; S, spinning stage; PP, prepupa stage; P, pupa stage; SP, semi-pupa; D, dead individual. CK, control. After 72 h of spining, 100% entered the pupal stage. AM group, HPPC model group. The low-melting point wax (55℃-60℃) blocked the spinning holes at wanderding stage, completely preventing the spining, and it was difficult to develop to adults. At 192 h, the percentage composition of pupa, semi-pupa and dead individuals was 4:44:52. mAM group, mild HPPC model group. The spinning hole was blocked for modeling after 24 hours of spinning, and some individuals were able to complete the development of the age. At 192 h, the percentage composition of pupa, semi-pupa and dead individuals was 42:26:32. The black arrow indicated the individual injection treatment. The mAM group was injected with endocrine hormone active substance 20E for rescue at 0 h or 24 h after modeling; The mAM and CK group were injected with siRNA-Gcm at 96 h (the time axis 120 h) after modeling; The AM group was injected with JAK inhibitor AG490 at 144 h after modeling. The blue arrow indicated the sampling survey time. (**B**) PPC level (n = 3). (**C-G**) Statistics of different types of blood cell density (n = 10). C, Pro; D, Gra; E, Pla; F, Oen; G, Sph. The ordinate represented the number of hemocytes per cubic millimeter of hemolymph. Data are shown as mean ± SEM. ns p>0.05,*p<0.05, **p<0.01 by Student’s T-test.

**Figure supplement 2.**
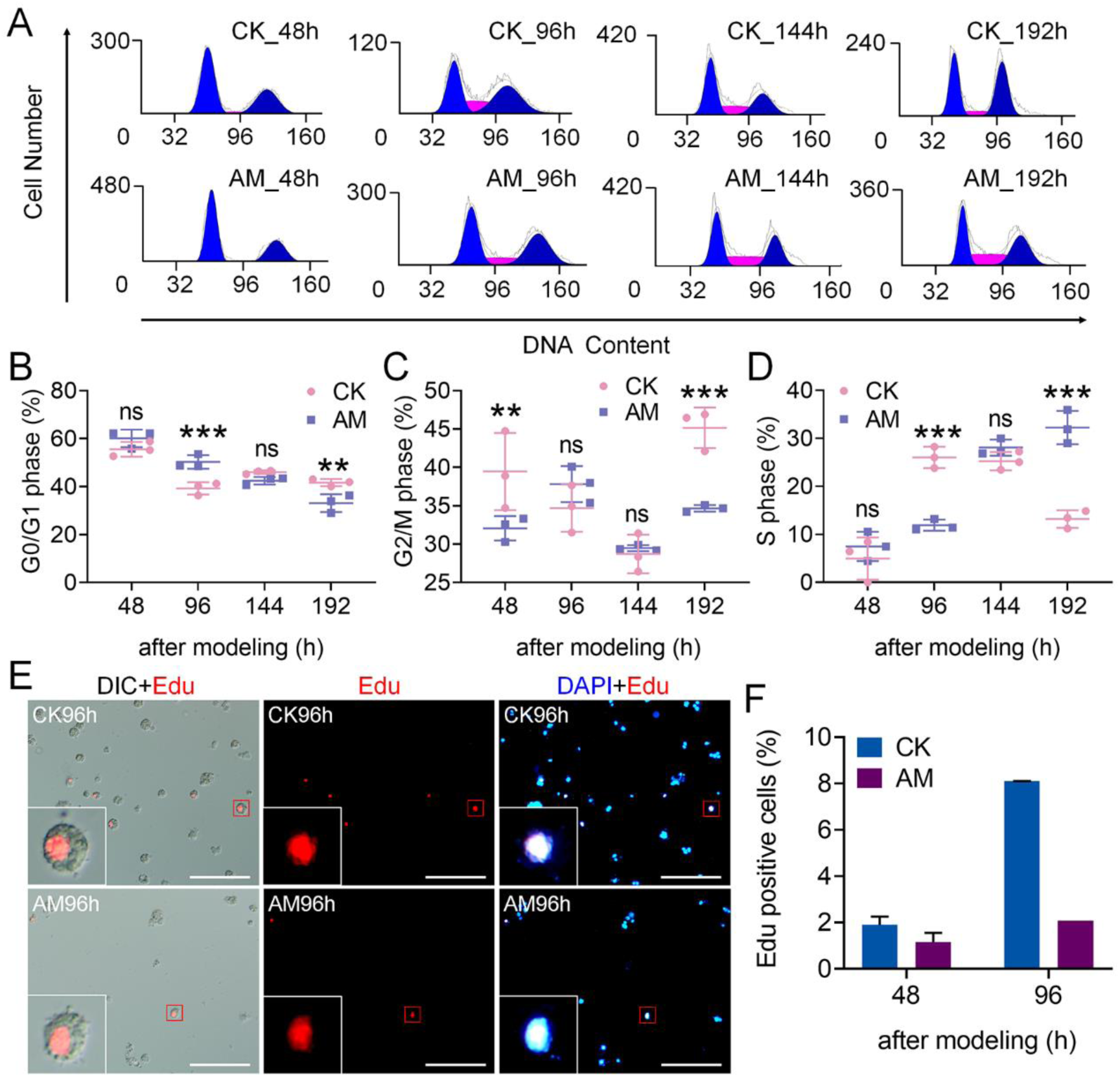
HPPC affected the cycle phase distribution of circulating hemocytes. Circulating hemocytes were investigated 48 h, 96 h, 144 h and 192 h after modeling. The cell cycle was divided into G0/G1 phase, S phase and G2/M phase. (**A**) The phase distribution of blood cell cycle were detected by PI-FCM (n = 3). (**B-D**) The percentage of blood cells in each phase (n = 3). **B**, G0/G1 phase, showing cells before DNA replication, at which time the cells did not proliferate; **C**, S phase, showing cells in DNA replication; **D**, G2/M phase, showing the cell division after DNA replication. Data are shown as mean ± SEM. ns p>0.05,*p<0.05, **p<0.01, and ***p<0.001 by Student’s T-test. (**E-F**) The second experiment of Edu staining. **E**, Fluorescence of Edu staining at 96 h after modeling. Scale bars 100 μm. **F**, Edu-positive rate of cells at 48 h and 96 h after modeling.

**Figure supplement 3.**
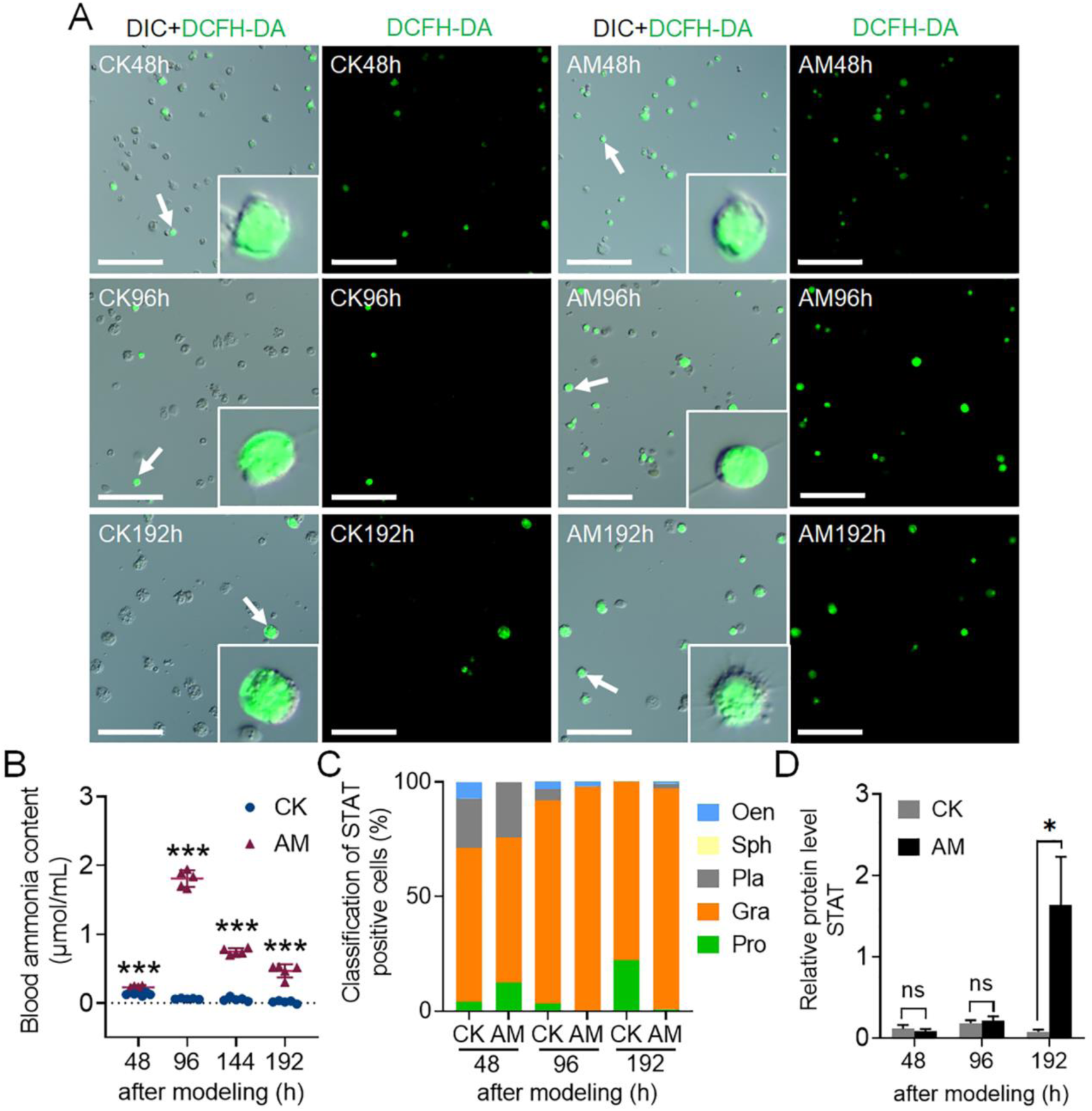
HPPC induced an increase in ROS levels in circulating hemocytes. (**A**) Fluorescence images of hemocytes by DCFH-DA staining at 48 h, 96 h, and 192 h after modeling. Scale bars 100 μm. (**B**) Changes of blood ammonia levels in hemolymph between 48 h and 192 h after modeling (n = 5). (**C**) Classification of STAT-positive blood cell composition. The STAT- positive rate of different types of hemocytes (%) = (number of STAT positive hemocytes of different types / total number of positive cells) × 100. (**D**) The relative expression level of STAT protein in blood cells was analyzed by western blotting at 48 h, 96 h and 192 h after modeling (n = 3). Referenced protein was β-Tubulin. In Figure S3B and Figure S3D, data are shown as mean ± SEM. ns p>0.05,*p<0.05 and ***p<0.001 by Student’s T-test.

**Figure supplement 4.**
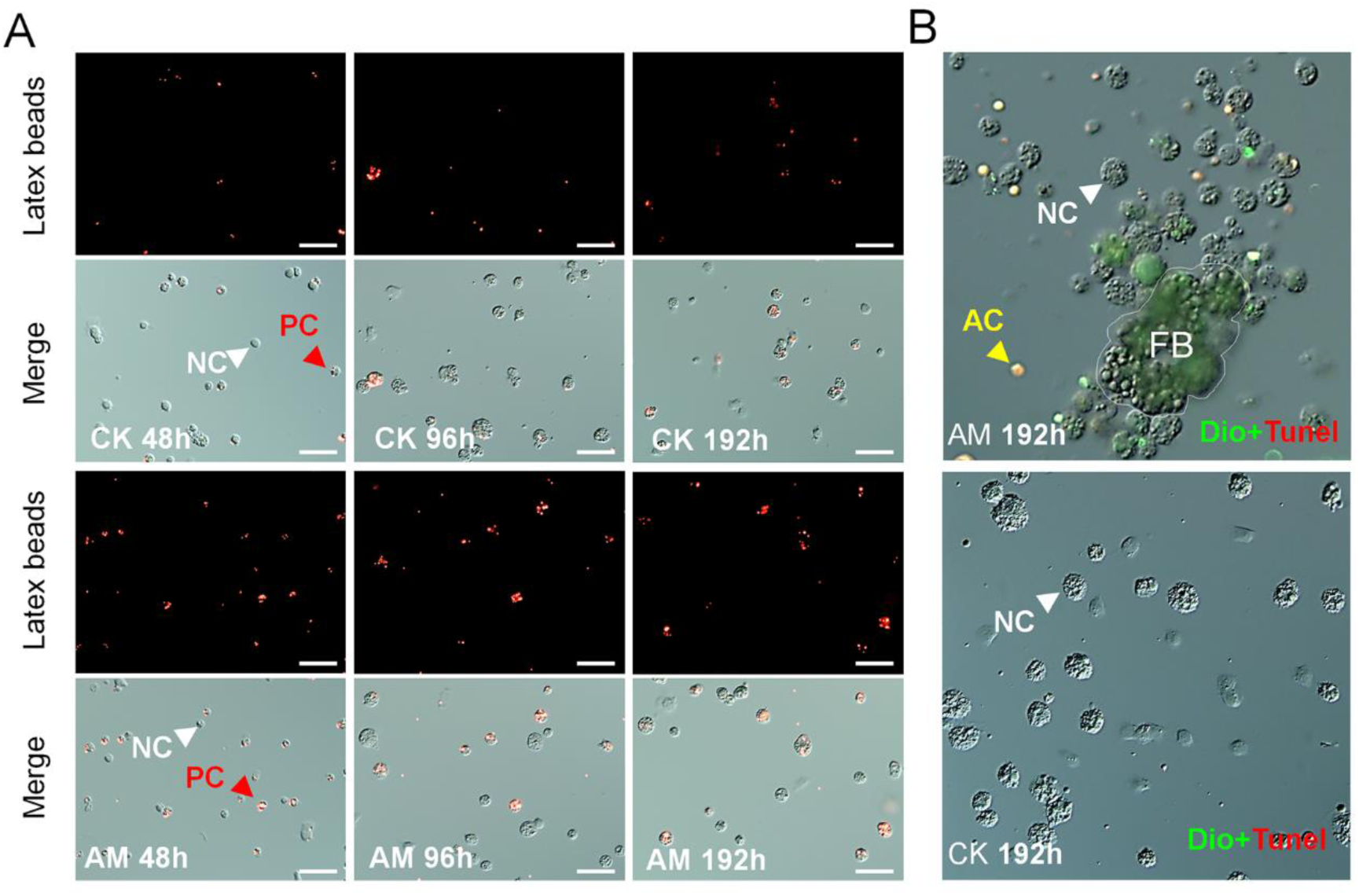
HPPC induced the increasing phagocytic capacity. (**A**) Fluorescent microspheres phagocytosed by hemocytes. Fluorescent microspheres were injected at 48 h, 96 h and 192 h after modeling, and the hemolymph was collected for microscopic observation after 4h in vivo incubation. PC, phagocytic blood cells; NC, normal blood cells. (**B**) Hemocytes phagocytized (fat body) tissue fragments 192 h after modeling. Dio, cell membrane staining; Tunel, apoptosis indicator; FB, fat body fragments; AC, apoptotic blood cells.

**Figure supplement 5.**
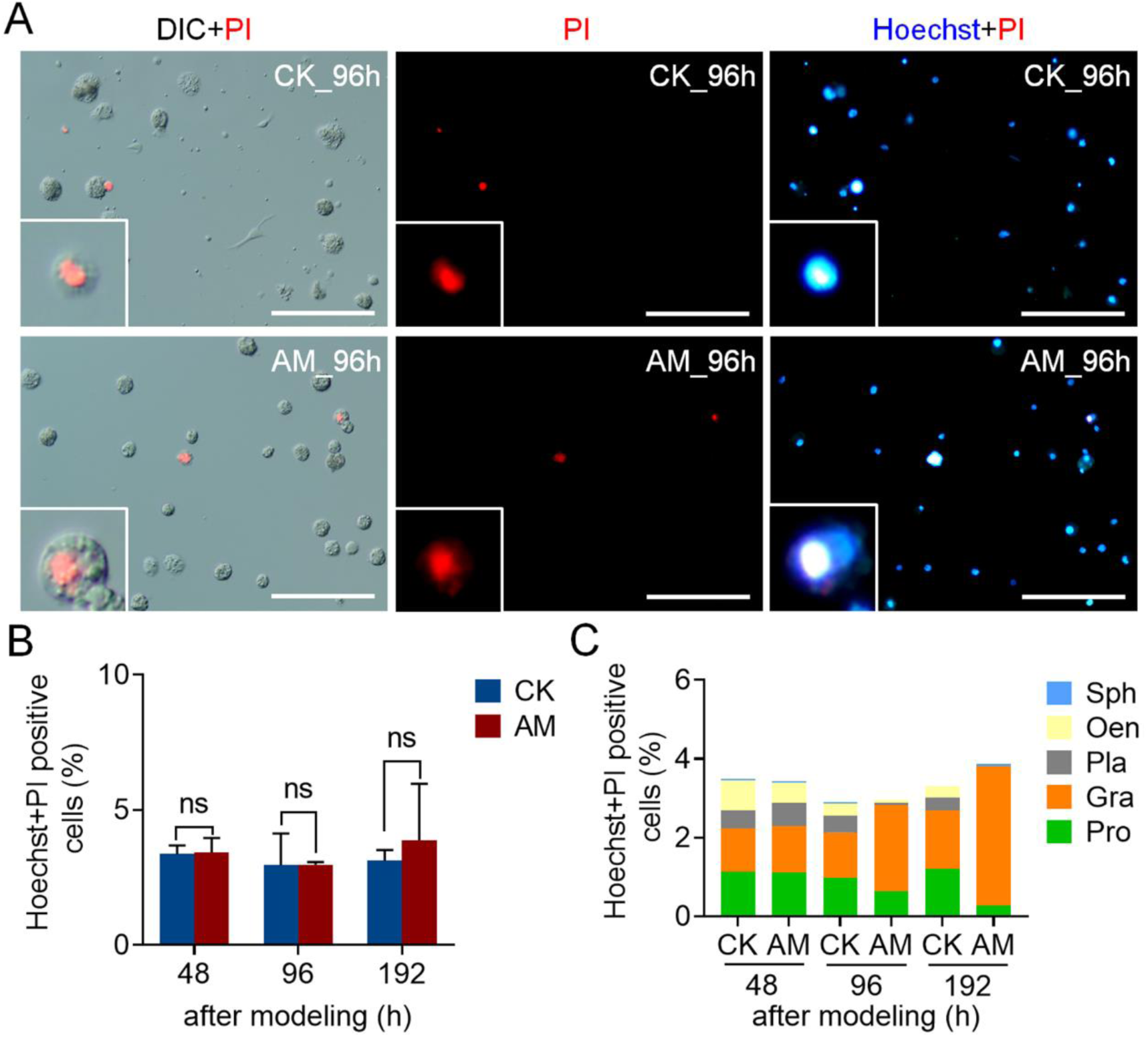
Investigation of blood cell necrosis level. Used Apoptosis and Necrosis Assay Kit (C1056, Beyotime, China) staining to detect necrosis of hemocytes. Took 60 μL blood sample, diluted 5-10 times (v/v) with HBSS, and climbed the slide for 5 min. After washing with PBS for 3 times, added 1000 μL of cell staining working solution, and incubated for 30 min in an ice bath in the dark. After washing with PBS for 3 times in the dark, observed the blue fluorescence (352/461 nm) and red fluorescence (488/630 nm). Hoechst+Propidium Iodide (PI) staining marked the necrotic hemocytes in the hemolymph at 48 h, 96 h and 192 h after modeling. DIC meaned cells in bright field, and Hoechst+PI double positive meaned necrotic cells. Positive rate (%) = (Hoechst+PI double positive cell number / total cell number) × 100. (**A**) 96 h Hoechst+PI staining fluorescence image after modeling. Scale bars 100 μm. (**B**) Hoechst+PI staining positive rate of hemocytes (n = 3). (**C**) Hoechst+PI staining positive rate of different types of hemocytes. Data are shown as mean ± SEM. ns p>0.05,*p<0.05, **p<0.01, and ***p<0.001 by Student’s T-test.

**Figure supplement 6.**
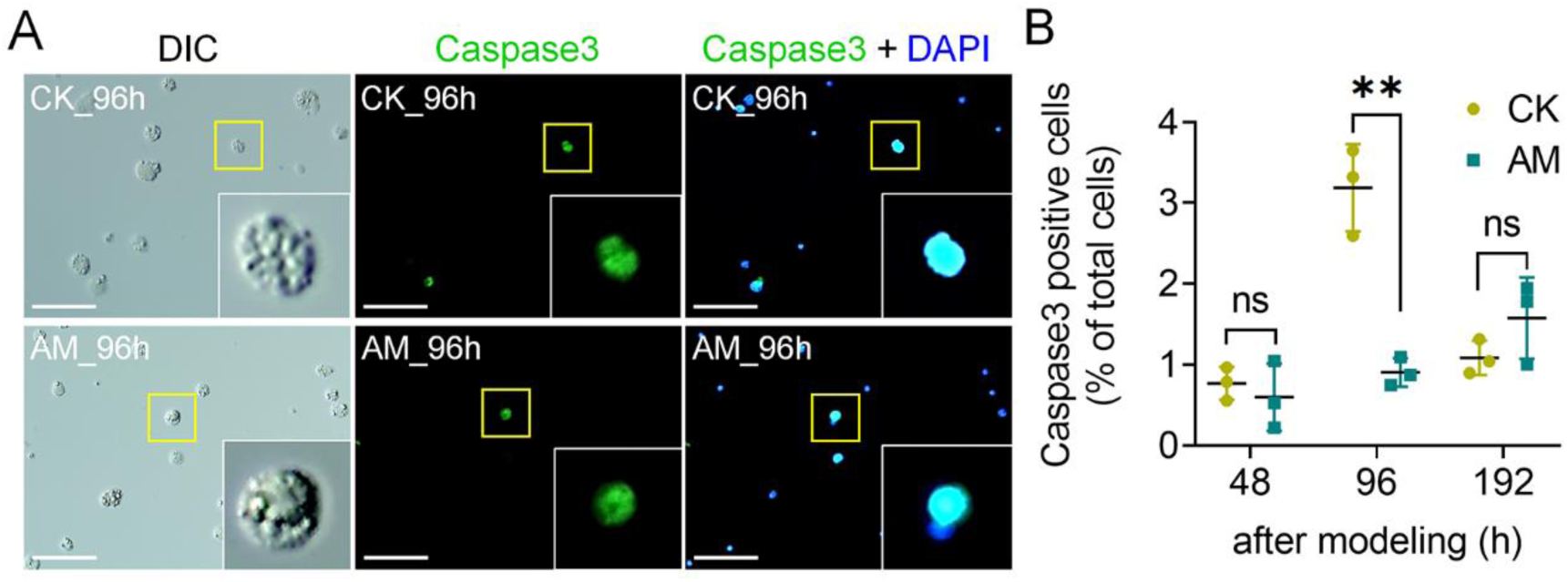
The expression level of Caspase-3 in blood cells was detected by immunofluorescence after modeling. Caspase-3 immunofluorescence staining marked the hemocytes expressing STAT protein in the hemolymph at 48 h, 96 h, and 192 h after modeling, and DAPI marked all cell nucleus (n = 3). (**A**) Caspase-3 immunostaining fluorescence image at 96 h after modeling. Scale bars 50 μm. (**B**) Caspase-3 immunofluorescence staining positive rate of hemocytes. Positive rate (%) = (STAT immunofluorescence positive cell number / total cell number) × 100. Data are shown as mean ± SEM. ns p>0.05, **p<0.01, and ***p<0.001 by Student’s T-test.

**Figure supplement 7.**
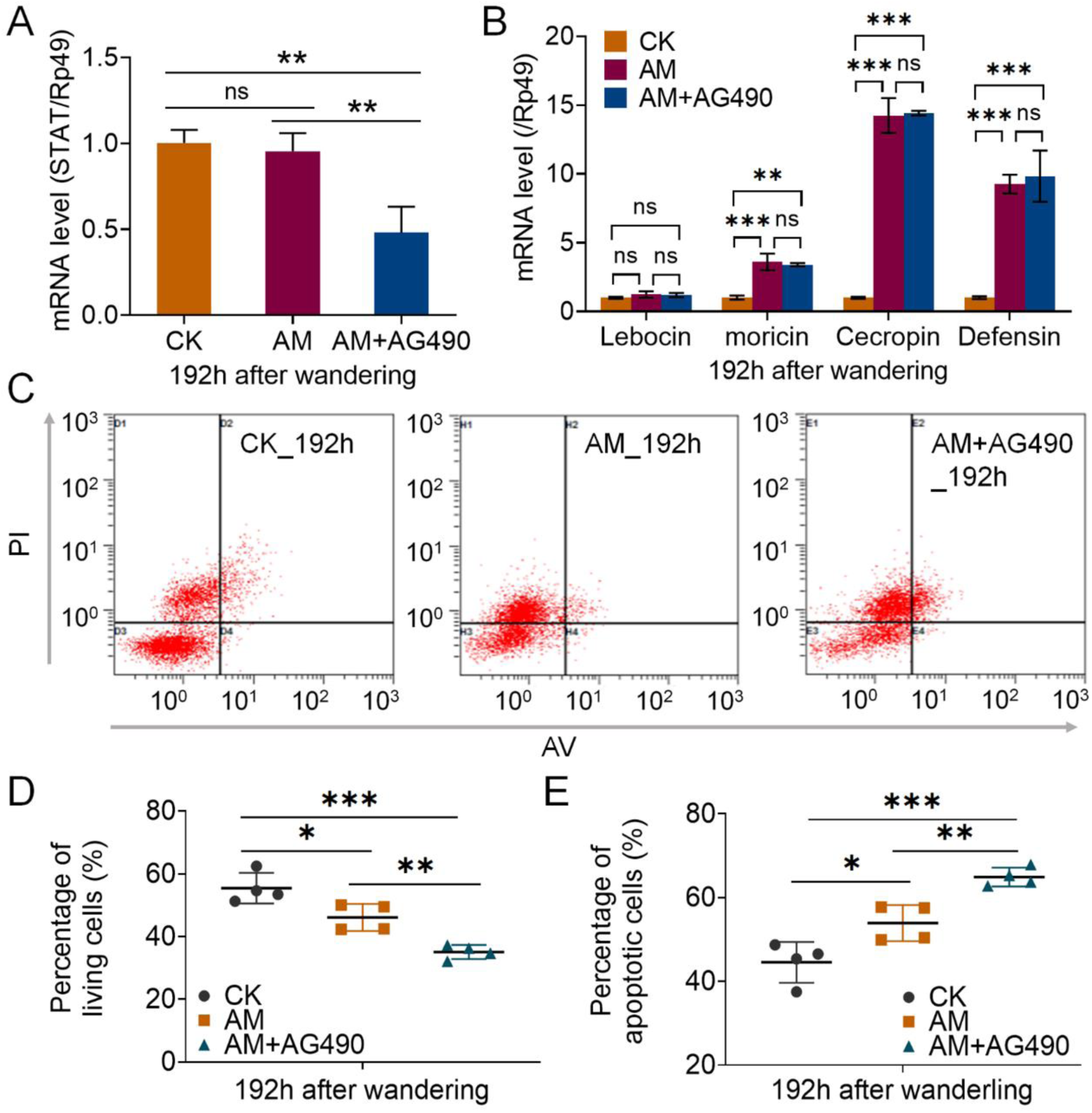
The effect of JAK inhibitor injection on the level of blood cell apoptosis. AM+AG490, AM group was injected with JAK inhibitor AG490 (per individual, 50 μmol/10 μL) at 24 hours after modeling, and the hemolymph was collected 168 h after injection. The mRNA levels of (**A**) STAT and (**B**) antimicrobial peptide gene in circulating hemocytes were detected by qPCR (n = 3). The reference gene was *Bombyx mori* Rp49. (**C**) Annexin V-FITC/PI - FCM results of blood cells. Negative AV and PI indicated living cells, while the others were apoptotic cells. The hemocytes in the AM group showed promotion of apoptosis after injection of AG490. (**D-E**) Annexin V-FITC/PI staining statistics (n = 4). **D**, the percentage of living cells. **E**, the percentage of apoptotic cells. Data are shown as mean ± SEM. ns p>0.05,*p<0.05, **p<0.01, and ***p<0.001 by Student’s T-test.

**Figure supplement 8.**
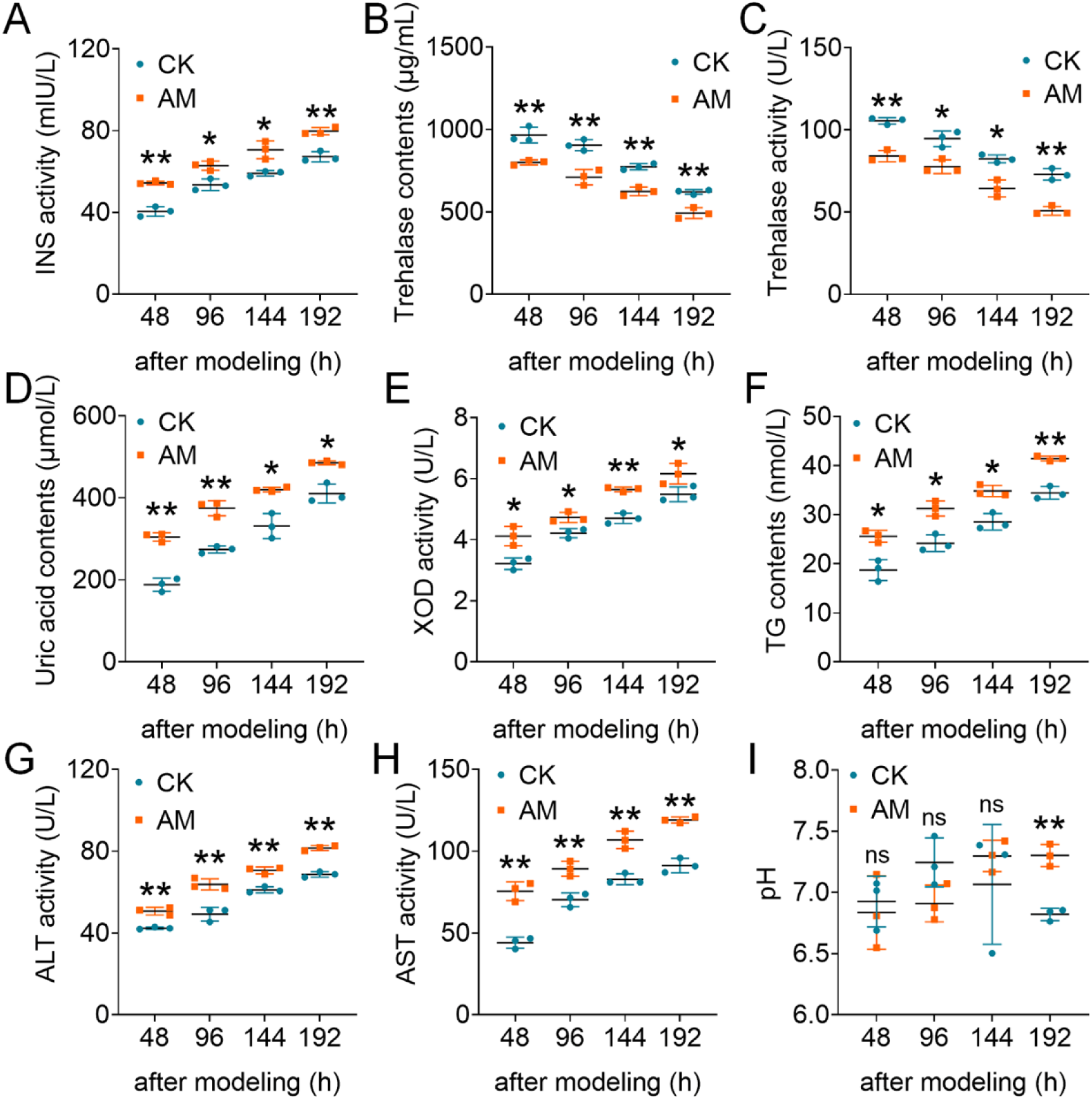
HPPC led to the synthesis and catabolism disorders of protein, sugar, lipid and other metabolites. The basic physiological parameters of hemolymph at 48 h, 96 h, 144 h and 192 h after modeling were determined by ELISA (n = 3). (**A**) INS activity; (**B**) Trehalase; (**C**) Trehalase activity; (**D**) Uric acid contents; (**E**) XOD activity; (**F**) TG contents; (**G**) ALT activity; (**H**) AST activity; (**I**) pH. Data are shown as mean ± SEM. The significant level of difference between AM and CK was marked as: ns p>0.05,*p<0.05, **p<0.01, and ***p<0.001 by Student’s T-test.

**Figure supplement 9.**
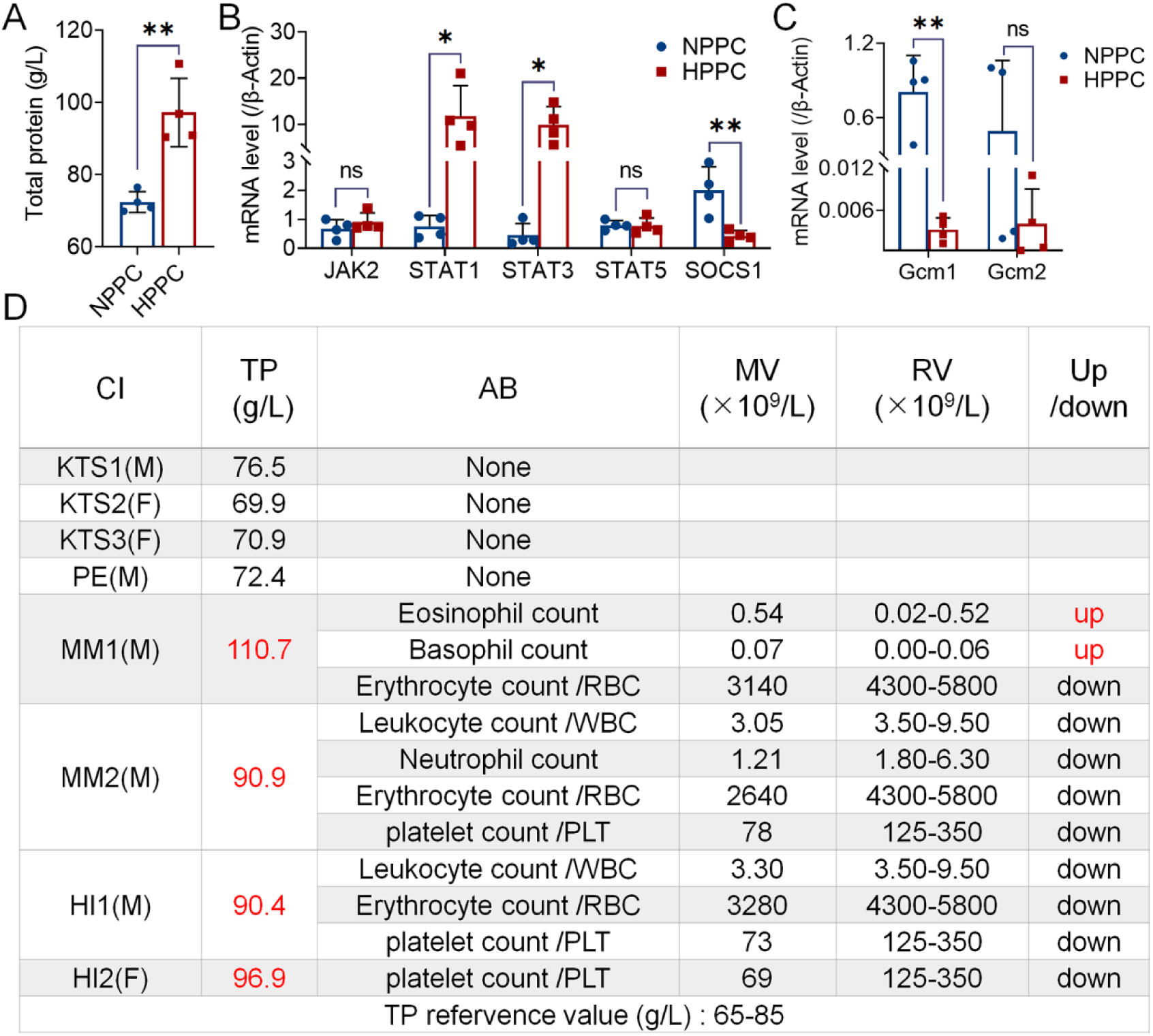
Investigation of JAK/STAT pathway and GCM gene transcription in peripheral blood of patients with HPPC (n = 4). NPPC, patients with normal total protein levels in clinical cases; HPPC, patients with total protein levels outside the normal range in clinical cases. (**A**) Total protein level, refervence value 65-85 g/L. (**B-C**) The transcription level of JAK/STAT pathway and Gcm-related genes in hemocytes investigated by qPCR. The internal reference gene was *Homo sapiens* β-Actin. Data are shown as mean ± SEM. ns p>0.05,*p<0.05, **p<0.01, and ***p<0.001 by Student’s T-test. (**D**) Clinical blood survey data. NPPC, normal PPC sample; HPPC, high PPC sample; M, male; F, female; CI, clinical impression; TP, total protein; AB, abnormal blood cell count; MV, measured value; RV, reference value; KTS, Kidney transplant status; PE, pleural effusion; MM, multiple myeloma; HI, hepatic impairment. RBC, Red blood cell count; WBC, White blood cell count.

## References

Abbas MN, Kausar S, Sun YX, Tian JW, Zhu BJ, Liu CL. 2018. Suppressor of cytokine signaling 6 can enhance epidermal growth factor receptor signaling pathway in Bombyx mori (Dazao). Developmental and comparative immunology 81:187–192. DOI: 10.1016/j.dci.2017.12.003, PMID: 29225004

Abuzaid AA, Aldahan MA, Helal MAA, Assiri AM, Alzahrani MH. 2020. Visceral leishmaniasis in Saudi Arabia: From hundreds of cases to zero. Acta tropica 212:105707. DOI: 10.1016/j.actatropica.2020.105707, PMID: 32950481

Adler BJ, Kaushansky K, Rubin CT. 2014. Obesity-driven disruption of haematopoiesis and the bone marrow niche. Nature reviews. Endocrinology 10:737–748. DOI: 10.1038/nrendo.2014.169, PMID: 25311396

Aguilar-Ballester M, Herrero-Cervera A, Vinué Á, Martínez-Hervás S, González-Navarro H. 2020. Impact of Cholesterol Metabolism in Immune Cell Function and Atherosclerosis. Nutrients 12:2021. DOI: 10.3390/nu12072021, PMID: 32645995

Bagchi S, He Y, Zhang H, Cao L, Van Rhijn I, Moody DB, Gudjonsson JE, Wang CR. 2017. CD1b-autoreactive T cells contribute to hyperlipidemia-induced skin inflammation in mice. Journal of clinical investigation 127:2339–2352. DOI: 10.1172/JCI92217, PMID: 28463230

Barrett TJ, Distel E, Murphy AJ, Hu J, Garshick MS, Ogando Y, Liu J, Vaisar T, Heinecke JW, Berger JS, Goldberg IJ, Fisher EA. 2019. Apolipoprotein AI) Promotes Atherosclerosis Regression in Diabetic Mice by Suppressing Myelopoiesis and Plaque Inflammation. Circulation 140:1170–1184. DOI: 10.1161/CIRCULATIONAHA.119.039476, PMID: 31567014

Bataillé L, Augé B, Ferjoux G, Haenlin M, Waltzer L. 2005. Resolving embryonic blood cell fate choice in Drosophila: interplay of GCM and RUNX factors. Development 132:4635–4644. DOI: 10.1242/dev.02034, PMID: 16176949

Bazzi W, Cattenoz PB, Delaporte C, Dasari V, Sakr R, Yuasa Y, Giangrande A. 2018. Embryonic hematopoiesis modulates the inflammatory response and larval hematopoiesis in Drosophila. eLife 7:e34890. DOI: 10.7554/eLife.34890, PMID: 29992900

Bergstedt J, Lingen C. Cirrhosis of the liver in two children with hyperproteinaemia. 1957. Acta paediatrica 46:185–190. DOI: 10.1111/j.1651-2227.1957.tb08643.x, PMID: 13424265

Boyle TE, Holowaychuk MK, Adams AK, Marks SL. 2011. Treatment of three cats with hyperviscosity syndrome and congestive heart failure using plasmapheresis. Journal of the American Animal Hospital Association 47:50–55. DOI: 10.5326/JAAHA-MS-5635, PMID: 21164170

Chang SH, Gumbel J, Luo S, Thomas TS, Sanfilippo KM, Luo J, Colditz GA, Carson KR. 2019. Post-MGUS Diagnosis Serum Monoclonal-Protein Velocity and the Progression of Monoclonal Gammopathy of Undetermined Significance to Multiple Myeloma. Cancer epidemiology, biomarkers & prevention 28:2055–2061. DOI: 10.1158/1055-9965.EPI-19-0132, PMID: 31501149

Chen XD, Wang YF, Wang YL, Li QY, Ma HY, Wang L, Sima YH, Xu SQ. 2018. Induced hyperproteinemia and its effects on the remodeling of fat bodies in silkworm, Bombyx mori. Frontiers in physiology 9:302. DOI: 10.3389/fphys.2018.00302, PMID: 29651251

Chistiakov DA, Grechko AV, Myasoedova VA, Melnichenko AA, Orekhov AN. 2018. The role of monocytosis and neutrophilia in atherosclerosis. Journal of cellular and molecular medicine 22:1366–1382. DOI: 10.1111/jcmm.13462, PMID: 29364567

Chong PSY, Chng WJ, de Mel S. 2019. STAT3: A Promising Therapeutic Target in Multiple Myeloma. Cancers (Basel) 11:731. DOI: 10.3390/cancers11050731, PMID: 31130718

Cui S, Lv X, Li W, Li Z, Liu H, Gao Y, Huang G. 2018. Folic acid modulates VPO1 DNA methylation levels and alleviates oxidative stress-induced apoptosis in vivo and in vitro. Redox biology 19:81–91. DOI: 10.1016/j.redox.2018.08.005, PMID: 30125807

Jaitin DA, Adlung L, Thaiss CA, Weiner A, Li B, Descamps H, Lundgren P, Bleriot C, Liu Z, Deczkowska A, Keren-Shaul H, David E, Zmora N, Eldar SM, Lubezky N, Shibolet O, Hill DA, Lazar MA, Colonna M, Ginhoux F, Shapiro H, Elinav E, Amit I. 2019. Lipid-Associated Macrophages Control Metabolic Homeostasis in a Trem2-Dependent Manner. Cell 178:686–698.e14. DOI: 10.1016/j.cell.2019.05.054, PMID: 31257031

Dodington DW, Desai HR, Woo M. JAK/STAT - Emerging Players in Metabolism. 2018. Trends in endocrinology and metabolism: TEM 29:55–65. DOI: 10.1016/j.tem.2017.11.001, PMID: 29191719

Fuchs Y, Steller H. 2015. Live to die another way: modes of programmed cell death and the signals emanating from dying cells. Nature reviews. Molecular cell biology 16:329–344. DOI: 10.1038/nrm3999, PMID: 25991373

Fujii M, Sato Y, Ohara N, Hashimoto K, Kobashi H, Koyama Y, Yoshino T. 2014. Systemic IgG4- related disease with extensive peripheral nerve involvement that progressed from localized IgG4- related lymphadenopathy: an autopsy case. Diagnostic pathology 9:41. DOI: 10.1186/1746-1596-9-41, PMID: 24559103

Galluzzi L, Vitale I, Aaronson SA, Abrams JM, Adam D, Agostinis P, Alnemri ES, Altucci L, Amelio I, Andrews DW, Annicchiarico-Petruzzelli M, Antonov AV, Arama E, Baehrecke EH, Barlev NA, Bazan NG, Bernassola F, Bertrand MJM, Bianchi K, Blagosklonny MV, Blomgren K, Borner C, Boya P, Brenner C, Campanella M, Candi E, Carmona-Gutierrez D, Cecconi F, Chan FK, Chandel NS, Cheng EH, Chipuk JE, Cidlowski JA, Ciechanover A, Cohen GM, Conrad M, Cubillos-Ruiz JR, Czabotar PE, D’Angiolella V, Dawson TM, Dawson VL, De Laurenzi V, De Maria R, Debatin KM, DeBerardinis RJ, Deshmukh M, Di Daniele N, Di Virgilio F, Dixit VM, Dixon SJ, Duckett CS, Dynlacht BD, El-Deiry WS, Elrod JW, Fimia GM, Fulda S, García-Sáez AJ, Garg AD, Garrido C, Gavathiotis E, Golstein P, Gottlieb E, Green DR, Greene LA, Gronemeyer H, Gross A, Hajnoczky G, Hardwick JM, Harris IS, Hengartner MO, Hetz C, Ichijo H, Jäättelä M, Joseph B, Jost PJ, Juin PP, Kaiser WJ, Karin M, Kaufmann T, Kepp O, Kimchi A, Kitsis RN, Klionsky DJ, Knight RA, Kumar S, Lee SW, Lemasters JJ, Levine B, Linkermann A, Lipton SA, Lockshin RA, López-Otín C, Lowe SW, Luedde T, Lugli E, MacFarlane M, Madeo F, Malewicz M, Malorni W, Manic G, Marine JC, Martin SJ, Martinou JC, Medema JP, Mehlen P, Meier P, Melino S, Miao EA, Molkentin JD, Moll UM, Muñoz-Pinedo C, Nagata S, Nuñez G, Oberst A, Oren M, Overholtzer M, Pagano M, Panaretakis T, Pasparakis M, Penninger JM, Pereira DM, Pervaiz S, Peter ME, Piacentini M, Pinton P, Prehn JHM, Puthalakath H, Rabinovich GA, Rehm M, Rizzuto R, Rodrigues CMP, Rubinsztein DC, Rudel T, Ryan KM, Sayan E, Scorrano L, Shao F, Shi Y, Silke J, Simon HU, Sistigu A, Stockwell BR, Strasser A, Szabadkai G, Tait SWG, Tang D, Tavernarakis N, Thorburn A, Tsujimoto Y, Turk B, Vanden Berghe T, Vandenabeele P, Vander Heiden MG, Villunger A, Virgin HW, Vousden KH, Vucic D, Wagner EF, Walczak H, Wallach D, Wang Y, Wells JA, Wood W, Yuan J, Zakeri Z, Zhivotovsky B, Zitvogel L, Melino G, Kroemer G. 2018. Molecular mechanisms of cell death: recommendations of the Nomenclature Committee on Cell Death. Cell death and differentiation 25:486–541. DOI: 10.1038/s41418-017-0012-4, PMID: 29362479

Gerin F, Ramazan DC, Baykan O, Sirikci O, Haklar G. 2014. Abnormal gel flotation in a patient with apperant pneumonia diagnosis: a case report. Biochemia medica 24:180–182. DOI: 10.11613/BM.2014.021, PMID: 24627728

Gu Q, Yang X, Lv J, Zhang J, Xia B, Kim JD, Wang R, Xiong F, Meng S, Clements TP, Tandon B, Wagner DS, Diaz MF, Wenzel PL, Miller YI, Traver D, Cooke JP, Li W, Zon LI, Chen K, Bai Y, Fang L. 2019. AIBP-mediated cholesterol efflux instructs hematopoietic stem and progenitor cell fate. Science 363:1085–1088. DOI: 10.1126/science.aav1749, PMID: 30705153

Hao Y, Jin LH. 2017. Dual role for Jumu in the control of hematopoietic progenitors in the Drosophila lymph gland. eLife 6:e25094. DOI: 10.7554/eLife.25094, PMID: 28350299

He Y , Xu X , Qiu J , Yin W, S Xu. 2019. Bombyx mori used as a fast detection model of liver melanization after a clinical drug – Acetaminophen exposure. Journal of Asia-Pacific Entomology. 23: 177–185. DOI: 10.1016/j.aspen.2019.11.009

Hu X, Zhang X, Wang J, Huang M, Xue R, Cao G, Gong C. 2015. Transcriptome analysis of BmN cells following over-expression of BmSTAT. Molecular genetics and genomics 290:2137–2146. DOI: 10.1007/s00438-015-1065-z, PMID: 25998838

Huang L, Liu D, Wang N, Ling S, Tang Y, Wu J, Hao L, Luo H, Hu X, Sheng L, Zhu L, Wang D, Luo Y, Shang Z, Xiao M, Mao X, Zhou K, Cao L, Dong L, Zheng X, Sui P, He J, Mo S, Yan J, Ao Q, Qiu L, Zhou H, Liu Q, Zhang H, Li J, Jin J, Fu L, Zhao W, Chen J, Du X, Qing G, Liu H, Liu X, Huang G, Ma D, Zhou J, Wang QF. 2018. Integrated genomic analysis identifies deregulated JAK/STAT-MYC-biosynthesis axis in aggressive NK-cell leukemia. Cell research 28:172–186. DOI: 10.1038/cr.2017.146, PMID: 29148541

Hussain A, Almenfi HF, Almehdewi AM, Hamza MS, Bhat MS, Vijayashankar NP. 2019. Laboratory Features of Newly Diagnosed Multiple Myeloma Patients. Cureus 11:e4716. DOI: 10.7759/cureus.4716, PMID: 31355076

Ishii K, Adachi T, Hamamoto H, Oonishi T, Kamimura M, Imamura K, Sekimizu K. 2013. Insect cytokine paralytic peptide activates innate immunity via nitric oxide production in the silkworm Bombyx mori. Developmental and comparative immunology 39:147–153. DOI: 10.1016/j.dci.2012.10.014, PMID: 23178406

Ishii K, Hamamoto H, Kamimura M, Nakamura Y, Noda H, Imamura K, Mita K, Sekimizu K. 2010. Insect cytokine paralytic peptide (PP) induces cellular and humoral immune responses in the silkworm Bombyx mori. The Journal of biological chemistry 285:28635–28642. DOI: 10.1074/jbc.M110.138446, PMID: 20622022

Khan IM, Pokharel Y, Dadu RT, Lewis DE, Hoogeveen RC, Wu H, Ballantyne CM. 2016. Postprandial Monocyte Activation in Individuals With Metabolic Syndrome. The Journal of clinical endocrinology and metabolism 101:4195–4204. DOI: 10.1210/jc.2016-2732, PMID: 27575945

Kitazawa A, Koda R, Yoshino A, Ueda Y, Takeda T. 2018. An IgA1-lambda-type monoclonal immunoglobulin deposition disease associated with membranous features in a patient with IgG4- related kidney disease: a case report. BMC nephrology 19:330. DOI: 10.1186/s12882-018-1133-9, PMID: 30458736

Kluck GEG, Wendt CHC, Imperio GED, Araujo MFC, Atella TC, da Rocha I, Miranda KR, Atella GC. 2019. Plasmodium Infection Induces Dyslipidemia and a Hepatic Lipogenic State in the Host through the Inhibition of the AMPK-ACC Pathway. Scientific reports 9:14695. DOI: 10.1038/s41598-019-51193-x, PMID: 31604978

Koranteng F, Cha N, Shin M, Shim J. 2020. The Role of Lozenge in Drosophila Hematopoiesis. Molecules and cells 43:114–120. DOI: 10.14348/molcells.2019.0249. PMID: 31992020

Kraakman MJ, Lee MK, Al-Sharea A, Dragoljevic D, Barrett TJ, Montenont E, Basu D, Heywood S, Kammoun HL, Flynn M, Whillas A, Hanssen NM, Febbraio MA, Westein E, Fisher EA, Chin- Dusting J, Cooper ME, Berger JS, Goldberg IJ, Nagareddy PR, Murphy AJ. 2017. Neutrophil- derived S100 calcium-binding proteins A8/A9 promote reticulated thrombocytosis and atherogenesis in diabetes. The Journal of clinical investigation 127:2133–2147. DOI: 10.1172/JCI92450, PMID: 28504650

Laforge M, Elbim C, Frère C, Hémadi M, Massaad C, Nuss P, Benoliel JJ, Becker C. 2020. Tissue damage from neutrophil-induced oxidative stress in COVID-19. Nature reviews Immunology 20:515–516. DOI: 10.1038/s41577-020-0407-1, PMID: 32728221

Lebestky T, Chang T, Hartenstein V, Banerjee U. 2000. Specification of Drosophila hematopoietic lineage by conserved transcription factors. Science 288:146–149. DOI: 10.1126/science.288.5463.146, PMID: 10753120

Li KL, Zhang YH, Xing R, Zhou YF, Chen XD, Wang H, Song B, Sima YH, He Y, Xu SQ. 2017. Different toxicity of cadmium telluride, silicon, and carbon nanomaterials against hemocytes in silkworm, Bombyx mori. RSC Advances 7: 50317–50327. DOI: 10.1039/C7RA09622D

Ling E, Shirai K, Kanekatsu R, Kiguchi K. 2003. Classification of larval circulating hemocytes of the silkworm, Bombyx mori, by acridine orange and propidium iodide staining. Histochemistry and cell biology 120:505–511. DOI: 10.1007/s00418-003-0592-6, PMID: 14610679

Ling E, Shirai K, Kanekatsu R, Kiguchi K. 2005. Hemocyte differentiation in the hematopoietic organs of the silkworm, Bombyx mori: prohemocytes have the function of phagocytosis. Cell and tissue research 320:535–543. DOI: 10.1007/s00441-004-1038-8, PMID: 15846518

Liu T, Xing R, Zhou YF, Zhang J, Su YY, Zhang KQ, He Y, Sima YH, Xu SQ. 2014. Hematopoiesis toxicity induced by CdTe quantum dots determined in an invertebrate model organism. Biomaterials 35:2942–2951. DOI: 10.1016/j.biomaterials.2013.12.007, PMID: 24411333

Matoušek V, Herold I, Holanová L, Balík M. 2018. A Rare Case of Severe Metabolic Alkalosis with Unusual Hyperproteinemia Treated with Continuous Renal Replacement Therapy and Regional Citrate Anticoagulation. Case reports in nephrology and dialysis 8:138–146. DOI: 10.1159/000491628, PMID: 30197902

Mattisson IY, Björkbacka H, Wigren M, Edsfeldt A, Melander O, Fredrikson GN, Bengtsson E, Gonçalves I, Orho-Melander M, Engström G, Almgren P, Nilsson J. 2017. Elevated Markers of Death Receptor-Activated Apoptosis are Associated with Increased Risk for Development of Diabetes and Cardiovascular Disease. EBioMedicine 26:187–197. DOI: 10.1016/j.ebiom.2017.11.023, PMID: 29208468

Nagareddy PR, Murphy AJ, Stirzaker RA, Hu Y, Yu S, Miller RG, Ramkhelawon B, Distel E, Westerterp M, Huang LS, Schmidt AM, Orchard TJ, Fisher EA, Tall AR, Goldberg IJ. 2013. Hyperglycemia promotes myelopoiesis and impairs the resolution of atherosclerosis. Cell metabolism 17:695–708. DOI: 10.1016/j.cmet.2013.04.001, PMID: 23663738

Nakahara Y, Kanamori Y, Kiuchi M, Kamimura M. 2010. Two hemocyte lineages exist in silkworm larval hematopoietic organ. PloS one 5:e11816. DOI: 10.1371/journal.pone.0011816, PMID: 20676370

Oliveira VDC, Mendes Junior AAV, Cavalcanti MCH, Madeira MF, Ferreira LC, Figueiredo FB, Campos MP, Nadal NV, Almosny NRP, Menezes RC. 2019. First description of parasite load and clinicopathological and anatomopathological changes in a dog naturally coinfected with Dioctophyme renale and Leishmania infantum in Brazil. Veterinary parasitology, regional studies and reports 18:100351. DOI: 10.1016/j.vprsr.2019.100351, PMID: 31796167

Qiu JF, Li X, Cui WZ, Liu XF, Tao H, Yang K, Dai TM, Sima YH, Xu SQ. 2019. Inhibition of period gene expression causes repression of cell cycle progression and cell growth in the Bombyx mori Cells. Frontiers in physiology 10:537. DOI: 10.3389/fphys.2019.00537, PMID: 31130878

Raivola J, Haikarainen T, Silvennoinen O. 2019. Characterization of JAK1 Pseudokinase Domain in Cytokine Signaling. Cancers (Basel) 12:78. DOI: 10.3390/cancers12010078, PMID: 31892268

Recio C, Lucy D, Purvis GSD, Iveson P, Zeboudj L, Iqbal AJ, Lin D, O’Callaghan C, Davison L, Griesbach E, Russell AJ, Wynne GM, Dib L, Monaco C, Greaves DR. 2018. Activation of the Immune-Metabolic Receptor GPR84 Enhances Inflammation and Phagocytosis in Macrophages. Frontiers in immunology 9:1419. DOI: 10.3389/fimmu.2018.01419, PMID: 29973940

Ren Z, Ahn JH, Liu H, Tsai YH, Bhanu NV, Koss B, Allison DF, Ma A, Storey AJ, Wang P, Mackintosh SG, Edmondson RD, Groen RWJ, Martens AC, Garcia BA, Tackett AJ, Jin J, Cai L, Zheng D, Wang GG. 2019. PHF19 promotes multiple myeloma tumorigenicity through PRC2 activation and broad H3K27me3 domain formation. Blood 134:1176–1189. DOI: 10.1182/blood.2019000578, PMID: 31383640

Riemer F, Kuehner KA, Ritz S, Sauter-Louis C, Hartmann K. 2016. Clinical and laboratory features of cats with feline infectious peritonitis--a retrospective study of 231 confirmed cases (2000- 2010). Journal of feline medicine and surgery 18:348–356. DOI: 10.1177/1098612X15586209, PMID: 26185109

Sant C, Thomas S, Wint C, Maharaj V, Kalloo N, Hosein A. 2020. A case of Pearsonema eggs in the urine sediment of a cat in Trinidad. Veterinary Parasitology- Regional Studies and Reports 22:100491. DOI: 10.1016/j.vprsr.2020.100491, PMID: 33308735

Sarrazy V, Viaud M, Westerterp M, Ivanov S, Giorgetti-Peraldi S, Guinamard R, Gautier EL, Thorp EB, De Vivo DC, Yvan-Charvet L. 2016. Disruption of Glut1 in Hematopoietic Stem Cells Prevents Myelopoiesis and Enhanced Glucose Flux in Atheromatous Plaques of ApoE(-/-) Mice. Circulation research 118:1062–1077. DOI: 10.1161/CIRCRESAHA.115.307599, PMID: 26926469

Silwal P, Kim JK, Kim YJ, Jo EK. 2020. Mitochondrial Reactive Oxygen Species: Double-Edged Weapon in Host Defense and Pathological Inflammation During Infection. Frontiers in immunology 11:1649. DOI: 10.3389/fimmu.2020.01649, PMID: 32922385

Simion V, Zhou H, Haemmig S, Pierce JB, Mendes S, Tesmenitsky Y, Pérez-Cremades D, Lee JF, Chen AF, Ronda N, Papotti B, Marto JA, Feinberg MW. 2020. A macrophage-specific lncRNA regulates apoptosis and atherosclerosis by tethering HuR in the nucleus. Nature communications 11:6135. DOI: 10.1038/s41467-020-19664-2, PMID: 33262333

Steinberger BA, Ford SM, Coleman TA. 2003. Intravenous immunoglobulin therapy results in post-infusional hyperproteinemia, increased serum viscosity, and pseudohyponatremia. American journal of hematology 73:97–100. DOI: 10.1002/ajh.10325, PMID: 12749010

Tabunoki H, Bono H, Ito K, Yokoyama T. 2016. Can the silkworm (Bombyx mori) be used as a human disease model? Drug discoveries & therapeutics 10:3–8. DOI:10.5582/ddt.2016.01011, PMID: 26853920

Tall AR, Yvan-Charvet L. 2015. Cholesterol, inflammation and innate immunity. Nature reviews. Immunology 15:104–116. DOI: 10.1038/nri3793, PMID: 25614320

Trébuchet G, Cattenoz PB, Zsámboki J, Mazaud D, Siekhaus DE, Fanto M, Giangrande A. 2019. The Repo Homeodomain Transcription Factor Suppresses Hematopoiesis in Drosophila and Preserves the Glial Fate. The Journal of neuroscience 39:238–255. DOI: 10.1523/JNEUROSCI.1059-18.2018, PMID: 30504274

Vion AC, Kheloufi M, Hammoutene A, Poisson J, Lasselin J, Devue C, Pic I, Dupont N, Busse J, Stark K, Lafaurie-Janvore J, Barakat AI, Loyer X, Souyri M, Viollet B, Julia P, Tedgui A, Codogno P, Boulanger CM, Rautou PE. 2017. Autophagy is required for endothelial cell alignment and atheroprotection under physiological blood flow. Proceedings of the National Academy of Sciences of the United States of America 114:E8675–E8684. DOI: 10.1073/pnas.1702223114, PMID: 28973855

Wagner R, Heni M, Tabák AG, Machann J, Schick F, Randrianarisoa E, Hrabě de Angelis M, Birkenfeld AL, Stefan N, Peter A, Häring HU, Fritsche A. 2021. Pathophysiology-based subphenotyping of individuals at elevated risk for type 2 diabetes. Nature medicine 27:49–57. DOI: 10.1038/s41591-020-1116-9, PMID: 33398163

Wang YF, Chen XD, Wang G, Li QY, Liang XY, Sima YH, Xu SQ. 2019. Influence of hyperproteinemia on reproductive development in an invertebrate model. International journal of biological sciences 15:2170–2181. DOI: 10.7150/ijbs.33310, PMID: 31592097

Wang YF, Wang G, Li JL, Qu YX, Liang XY, Chen XD, Sima YH, Xu SQ. 2021. Influence of Hyperproteinemia on Insect Innate Immune Function of the Circulatory System in Bombyx mori. Biology (Basel) 10:112. DOI: 10.3390/biology10020112, PMID: 33546519

West AP, Brodsky IE, Rahner C, Woo DK, Erdjument-Bromage H, Tempst P, Walsh MC, Choi Y, Shadel GS, Ghosh S. 2011. TLR signalling augments macrophage bactericidal activity through mitochondrial ROS. Nature 472:476–480. DOI: 10.1038/nature09973, PMID: 21525932

Xi H, Zhang Y, Xu Y, Yang WY, Jiang X, Sha X, Cheng X, Wang J, Qin X, Yu J, Ji Y, Yang X, Wang H. 2016. Caspase-1 Inflammasome Activation Mediates Homocysteine-Induced Pyrop- Apoptosis in Endothelial Cells. Circulation research 118:1525–1539. DOI: 10.1161/CIRCRESAHA.116.308501, PMID, 27006445

Xu M, Wang X, Tan J, Zhang K, Guan X, Patterson LH, Ding H, Cui H. 2015. A novel Lozenge gene in silkworm, Bombyx mori regulates the melanization response of hemolymph. Developmental and comparative immunology 53:191–198. DOI: 10.1016/j.dci.2015.07.001, PMID, 26164197

Zhang M, Zhu H, Ding Y, Liu Z, Cai Z, Zou MH. 2017. AMP-activated protein kinase α1 promotes atherogenesis by increasing monocyte-to-macrophage differentiation. The Journal of biological chemistry 292:7888–7903. DOI: 10.1074/jbc.M117.779447, PMID: 28330873

Zhao H, Li Y, He L, Pu W, Yu W, Li Y, Wu YT, Xu C, Wei Y, Ding Q, Song BL, Huang H, Zhou B. 2020. In Vivo AAV-CRISPR/Cas9-Mediated Gene Editing Ameliorates Atherosclerosis in Familial Hypercholesterolemia. Circulation 141:67–79. DOI: 10.1161/CIRCULATIONAHA.119.042476, PMID: 31779484

